# Global Trends and Hotspots in Grassland Root Functional Traits Research (2000–2025)

**DOI:** 10.64898/2026.01.31.702986

**Authors:** Yongxue Feng, Yang Gao, Junqin Li, Yuting Yang, Jing Yin, Shu Yang, Zhihua Jiang, Xiangtao Wang, Puchang Wang, Lili Zhao

**Affiliations:** School of Life Sciences, Guizhou Normal University, Guiyang, Guizhou 550025, China; School of Karst Science, Guizhou Normal University, Guiyang, Guizhou 550025, China; College of Animal Science, Guizhou University, Guiyang, Guizhou 550025, China

**Keywords:** Grassland, Root Functional Traits, Bibliometrics, Soil

## Abstract

Grasslands, as one of the most important terrestrial ecosystems globally, have root functional traits that serve as key indicators of plant responses to environmental changes and hold significant ecological importance. To reveal the current status, research hotspots, and frontier trends in the field of grassland root functional traits, this study analyzed relevant literature from the Web of Science Core Collection database between 2000 and 2025. It employs bibliometric methods and utilizes visualization tools such as CiteSpace for a systematic analysis. The results indicate that research in this field has been continuously increasing since 2000, reflecting a growing research interest. China, the United States, and Germany are the leading countries in terms of publication output. However, collaboration networks among authors, institutions, and countries are still not tight enough to form a truly global cooperative network. Co-occurrence analysis of keywords and literature clustering reveal that the research hotspots in this field are mainly concentrated in six directions: multidimensional characteristics of root functional traits, interactions between root functional traits and climate change, synergistic effects of root functional traits and soil microorganisms, responses of root functional traits to land-use changes, coupling of root functional traits with ecosystem functions, and applications of root functional traits in agriculture and ecological restoration. Future research should focus on promoting innovation and standardization of research methods, conducting long-term monitoring, deeply exploring the mechanisms of root–microbe interactions, implementing cross-scale integrative research and model construction, and building international collaborative networks.

## Introduction

Plant roots play a crucial role in supporting the plant body, absorbing nutrients and water, and are the main organs sustaining growth and development of the belowground parts of vegetation. The root system and its symbionts are core components in maintaining multiple ecosystem functions[1, 2]. Root functional traits are important indicators of plant responses to environmental changes and of their impacts on ecosystem processes[3, 4]. “Functional traits” refer to key plant traits that significantly influence ecosystem functions and reflect plant responses to environmental changes[5]. These traits capture the plastic morphological and physiological changes that plants exhibit to adapt to their environment, revealing fundamental functional characteristics of plants[6]. In macro-scale root ecology, the databases FRED, TRY-root, and FungalRoot serve as essential resources for macro-analytical research. FRED 3.0 contains over 150 000 standardized observations of fine-root traits, enabling robust assessments of trait-environment relationships across ecosystems. TRY-root integrates multi-source datasets, comprising trait records for more than 6.24 million plant individuals, and therefore facilitates inter-regional comparisons of root strategies[7]. FungalRoot collates mycorrhizal association data for >14 000 plant species, providing the empirical basis for elucidating global distribution patterns and ecological drivers of mycorrhizal symbioses. Collectively, these databases deliver harmonized, high-quality data that underpin multi-scale macro-analyses in root ecology[8].Table 1 presents a classification of major root functional traits (based on the framework proposed by Bardgett et al. 2014) including the definition, measurement method, and ecological significance of each trait. Among these traits, specific root length (SRL) is a key indicator of plant resource acquisition efficiency and reflects plant ecological adaptation strategies[9]. Root diameter (D) is an important root morphological parameter that directly affects the root system’s ability to absorb water and nutrients[10]. Studies have shown that larger root diameters are often positively correlated with higher mycorrhizal fungal colonization rates, thereby enhancing the plant’s efficiency in acquiring soil nutrients[11, 12]. Root tissue density (RTD) characterizes the tolerance of plant roots to environmental stress and is closely related to nutrient storage capacity and stress resistance, making it an important indicator of plant resource allocation strategies[13]. Root exudates (such as organic acids, sugars, and amino acids) play a key role in regulating soil nutrient cycling[14]. These compounds significantly enhance the availability of carbon (C), nitrogen (N), phosphorus (P), and mineral elements (e.g. Ca, Mg) in the soil by stimulating microbial metabolism or through abiotic chemical processes[15–17]. Nitrogen, a key component of amino acids and enzymes, directly influences root metabolism and respiratory efficiency via root nitrogen content (RNC). Traditionally, root respiration was viewed as basal metabolism sustaining cell survival (e.g., basal mitochondrial activity)[18]. Bloom et al. (1992) challenged this, showing root respiration is dynamically coupled with nitrogen uptake and assimilation rather than a constant basal process[19]. Higher RNC typically links to more active root respiration, providing extra energy for nutrient uptake and potentially enhancing assimilation rates[20–23]. Moreover, fine root turnover is a crucial part of ecosystem carbon and nitrogen cycling: through the decay and decomposition of fine roots, nutrients sequestered by plants are released back into the soil, maintaining ecosystem material cycling and energy flow[24].

**Table 1:**
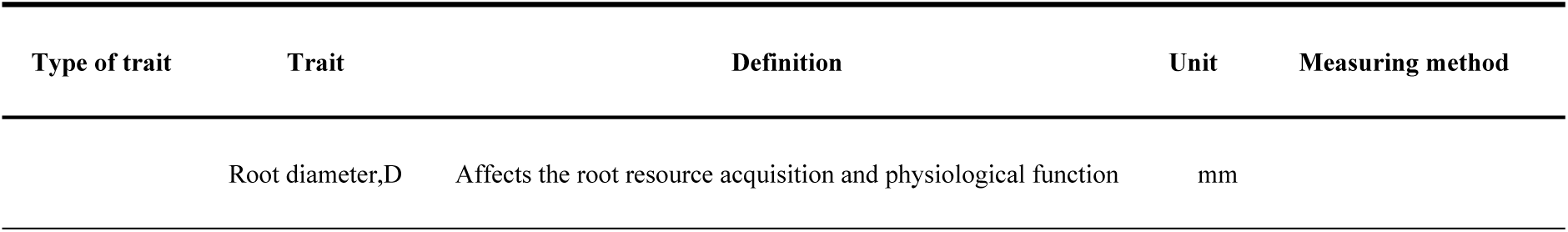
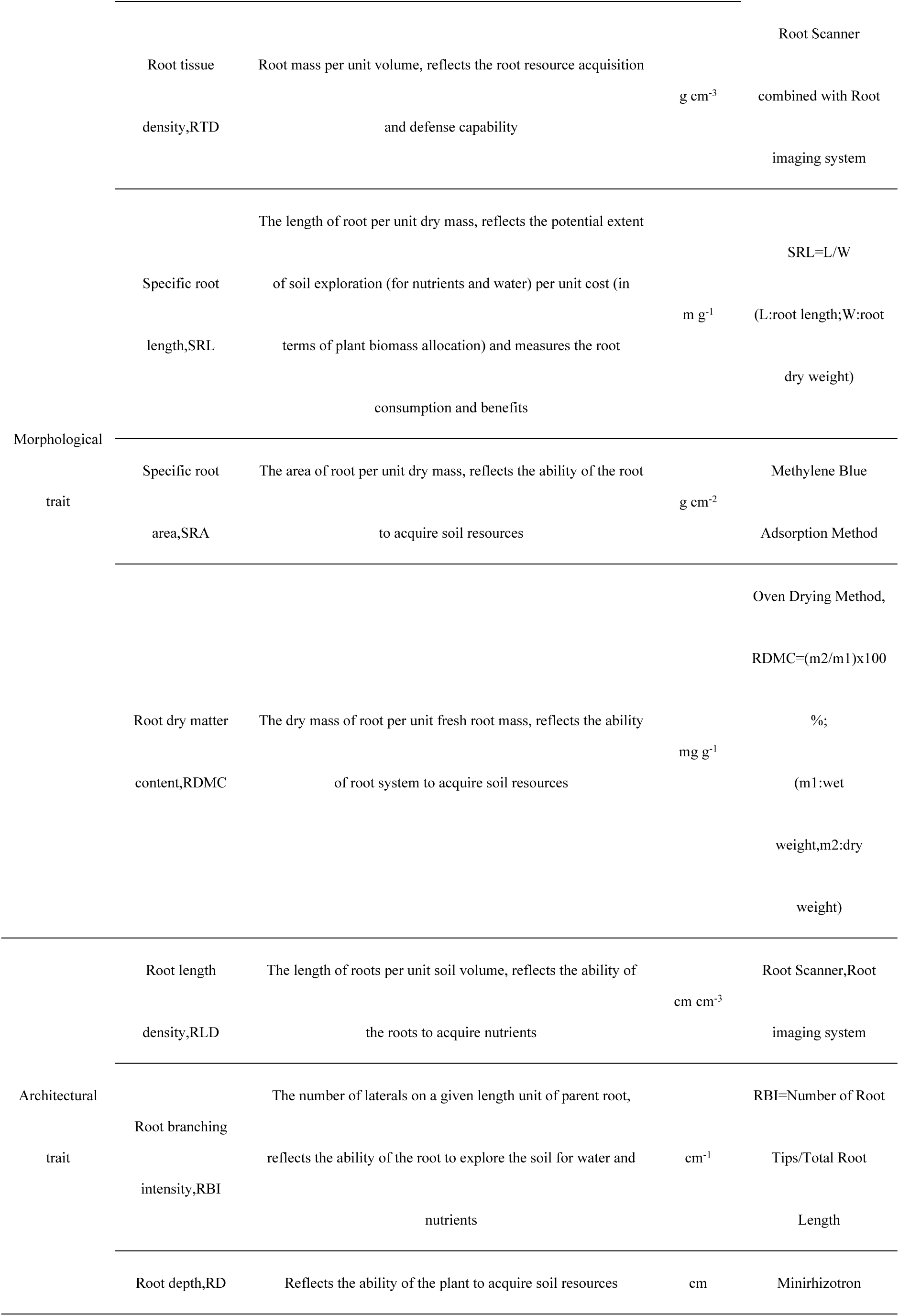
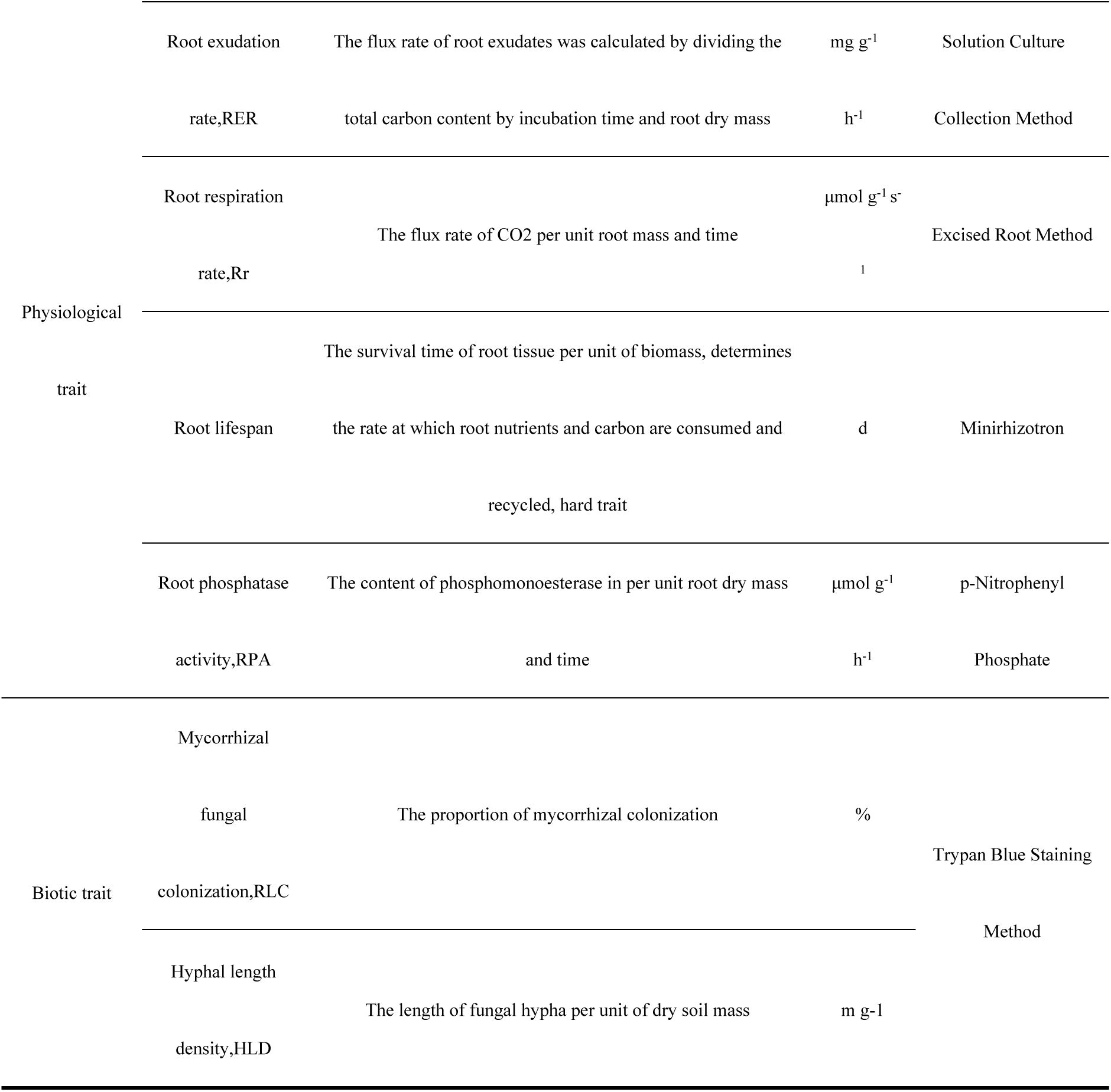
Classification, Units, Definitions, and Measurement Methods of Root Functional.

In nature, the total amount of resources available to plants is limited. However, plants have evolved strategies to cope with this limitation. Plants optimize resource acquisition through three primary strategies[25]: (1) Plastic adjustment of root architecture, which involves expanding the exploration range of soil resources by increasing root length or branching density[26, 27]; (2) Formation of mycorrhizal symbiotic networks, which enables absorption of nutrients beyond the immediate depletion zone of the rhizosphere via fungal hyphae[28, 29]; (3) Chemical regulation via root exudates, which enhances nutrient availability in the rhizosphere by releasing organic acids or enzymes[30, 31]. These strategies optimize the allocation of functional traits spatiotemporally, forming a trade-off mechanism between “resource acquisition” and “resource conservation”[32, 33]. Specifically, increased investment in certain root functional traits leads to reduced investment in others, manifesting as an enhanced advantage of some traits at the expense of others. This trade-off extends to the plant’s overall strategy along the “fast–slow” plant economics spectrum, which spans from rapid resource acquisition with high productivity to resource conservation with long lifespan[34]. In this context, the trade-off between root nutrient acquisition traits and conservation traits is conceptualized as the Root Economics Spectrum (RES)[35, 36]. The RES hypothesis posits that plant root traits are distributed along a gradient from acquisitive strategies (high resource uptake efficiency, short tissue lifespan) to conservative strategies (low uptake efficiency, long tissue lifespan)[37, 38]. However, whether a universal RES exists (comparable to the well-established Leaf Economics Spectrum) remains debated. For instance, [39]analyzed fine-root traits of 74 herbaceous and dwarf shrub species across Mediterranean, temperate, and tropical communities, and found that an acquisitive trait combination—high SRL, high root N content, and high root respiration rate—aligns with a carbon economy trade-off mechanism. This finding has been validated by subsequent studies on herbaceous plants[40, 41]. Conversely, a comprehensive review by [42]concluded that root trait variation is multidimensional, with no single dominant spectrum explaining all variation. Similarly, [43] showed that interactions between root functional traits and aboveground nitrogen retention mechanisms in grasslands deviate from predictions of the simple RES model. Further research indicates that the RES theory faces multiple challenges: (1) Root traits are driven by multiple factors (e.g. soil heterogeneity and mycorrhizal symbiosis) and may exhibit co-evolutionary coordination rather than strict trade-offs; (2) The divergence in root construction strategies between woody and herbaceous plants complicates applying RES universally across life forms; (3) Lack of standardized in situ observation techniques for three-dimensional root structural parameters (such as cortex thickness) limits data comparability; (4) Microbial interactions (e.g. mycorrhizal-mediated nutrient uptake pathways) are difficult to incorporate into the traditional RES framework[34, 42, 44].

Grasslands are under severe threat from ongoing degradation, undermining their capacity to support biodiversity, ecosystem services and human well-being[45]. In recent years, research on grassland aboveground components has been relatively extensive, whereas attention to the functional traits of belowground root systems has been comparatively insufficient. Roots support plant growth by absorbing and storing water and nutrients, prevent soil erosion, and enhance ecosystem stability and resilience. The diversity of root functional traits not only helps increase grassland biodiversity and productivity but also promotes soil health and nutrient cycling. Among these traits, the symbiosis between arbuscular mycorrhizal fungi (AMF) and roots plays a key role in nutrient uptake and soil structure maintenance[46]. Additionally, adaptive changes in roots help plants cope with environmental stresses such as drought and nutrient deficiency, providing theoretical support for grassland ecological restoration efforts[47, 48]. Understanding root responses to management practices (such as grazing and fertilization) is crucial for developing scientific grassland management strategies, breeding plant varieties adapted to changing conditions, and optimizing the sustainable use of grasslands[49]. Bibliometrics—using quantitative methods to analyze published literature—can systematically and accurately reveal the current status, developmental trajectory, and research hotspots of a given field[50]. To date, there has been relatively little scientometric research on grassland root functional traits. Therefore, this study applies scientometric approaches, in combination with tools such as VOSviewer, CiteSpace, and R, to analyze the knowledge structure and trends in the field of grassland root functional traits.

## 1. Data Sources and Research Methods

### 1.1 Data Sources and Screening

This study utilized the Web of Science Core Collection database as the data source, employing an advanced search strategy to retrieve relevant literature on the topic. The search query included the following English keyword groups: “functional traits of roots” or “root functional” or “root traits” or “belowground traits” or “root” or “rhizome” or “fine root” or “specific root length” or “root diameter” or “root tissue density” or “root nitrogen content” or “root economic spectrum” or “coarse root,” in combination with grassland-related terms such as “grassland” or “steppe” or “meadow” or “savannah” or “prairie” or “pampas” or “pasture.” Document types were limited to Article or Review, and the time span was 2000–2025. The search was conducted on November 11, 2025. The initial search yielded 2,747 documents. After manual screening to exclude documents not highly relevant to the topic or lacking author information, a total of 2008 documents were included for analysis.The specific retrieval and screening process is illustrated in Figure 1.

**Figure 1:**
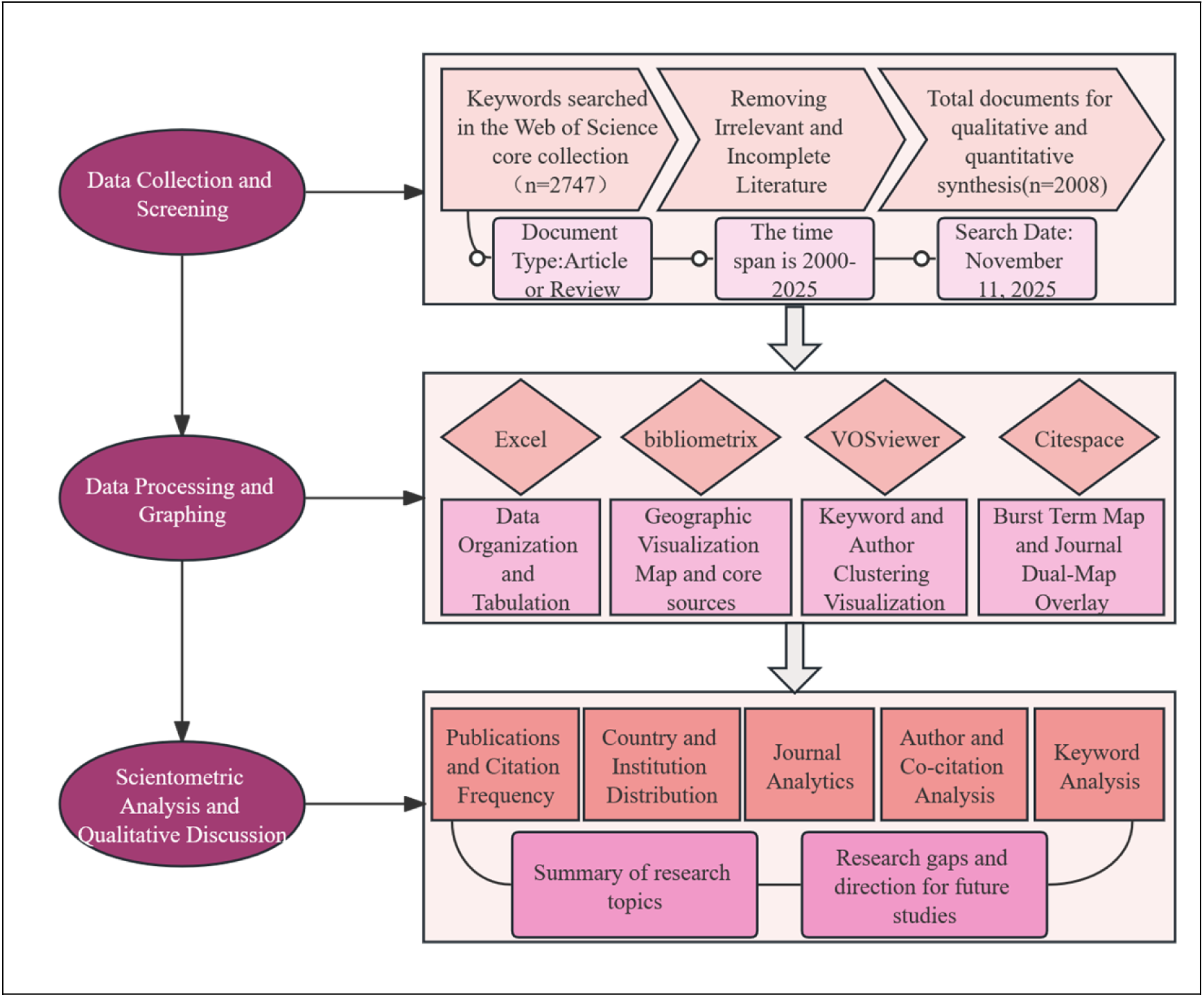
Flowchart of the Research Process on Grassland Root Functional Traits

### 1.2 Data Visualization and Analysis

Bibliometric analysis tools such as VOSviewer and the Bibliometrix R package were used to perform visualization and quantitative analysis of the grassland root functional traits research field. Key aspects analyzed included the most productive authors, research institutions, international cooperation networks, and keyword co-occurrence networks. In the author and country collaboration network maps, the size of each node represents the number of publications (and in the keyword co-occurrence map, node size represents keyword occurrence frequency)[51]. The lines connecting nodes indicate collaboration relationships between authors or countries (with thicker lines signifying closer collaboration)[52]; in the keyword co-occurrence map, the thickness of lines between keyword nodes indicates how frequently those keywords co-occur in the literature[53]. Clustering analysis and timeline visualizations (e.g., CiteSpace burst detection and temporal trend analyses) were also applied to identify research hotspots and emerging trends over time.

## 2. Results Analysis

### 2.1 Analysis of Publication Output and Highly Cited Documents

Conducting productivity analysis in a research field helps in understanding the dynamics and emerging trends within that field.Figure 2a illustrates the dynamic development of this research field from 2000 to 2025, characterized by a consistent upward trend in both annual and cumulative publications. The annual number of publications has increased from 31 in 2000 to 1,635 in 2023, with a predicted 1,870 publications in 2024. The red trend line fits the data well (R2= 0.94), confirming the robust growth trajectory. Notably, the growth accelerated after 2010, as evidenced by the steeper trend line. The cumulative publications curve demonstrates the rapid accumulation of research output, expected to reach approximately 3,760 by 2024. The confidence and prediction bands provide statistical support for the model’s reliability, indicating that the field has progressed from its initial phase to a mature, high-output stage. Figure 2b presents a joinpoint regression analysis identifying three significant turning points (2011, 2014, 2023) that delineate four distinct growth phases. The slopes for these periods are 2.89(2000-2011), 6.82(2011-2014), 2.69(2014-2023), and 15.43(2023-2025), respectively. The slope increase from 2.89 to 6.82 in 2011-2014 represents the first acceleration phase, likely driven by key methodological advances or conceptual breakthroughs. After a period of stable growth(2014-2023), the slope surged to 15.43 in 2023-2025, indicating the field is currently experiencing unprecedented expansion, possibly reflecting emerging research hotspots or increased interdisciplinary integration. Figure 2c reveals the evolution of high-impact research topics through citation analysis. Early seminal works focused on fundamental processes such as root turnover and rhizosphere effects, establishing core concepts in the field. The research focus subsequently shifted toward broader ecological applications, including soil carbon dynamics and plant CO₂ responses. Recent high-impact publications in prestigious journals like Nature Reviews Earth & Environment and Nature Communications address cutting-edge themes such as root economics spectrum, root and mycorrhizal nutrient acquisition strategies, and carbon and nitrogen cycling. This thematic progression from fundamental mechanisms to system-level applications demonstrates the field’s maturation and its increasing relevance to global challenges in ecosystem ecology and environmental science.

**Figure 2:**
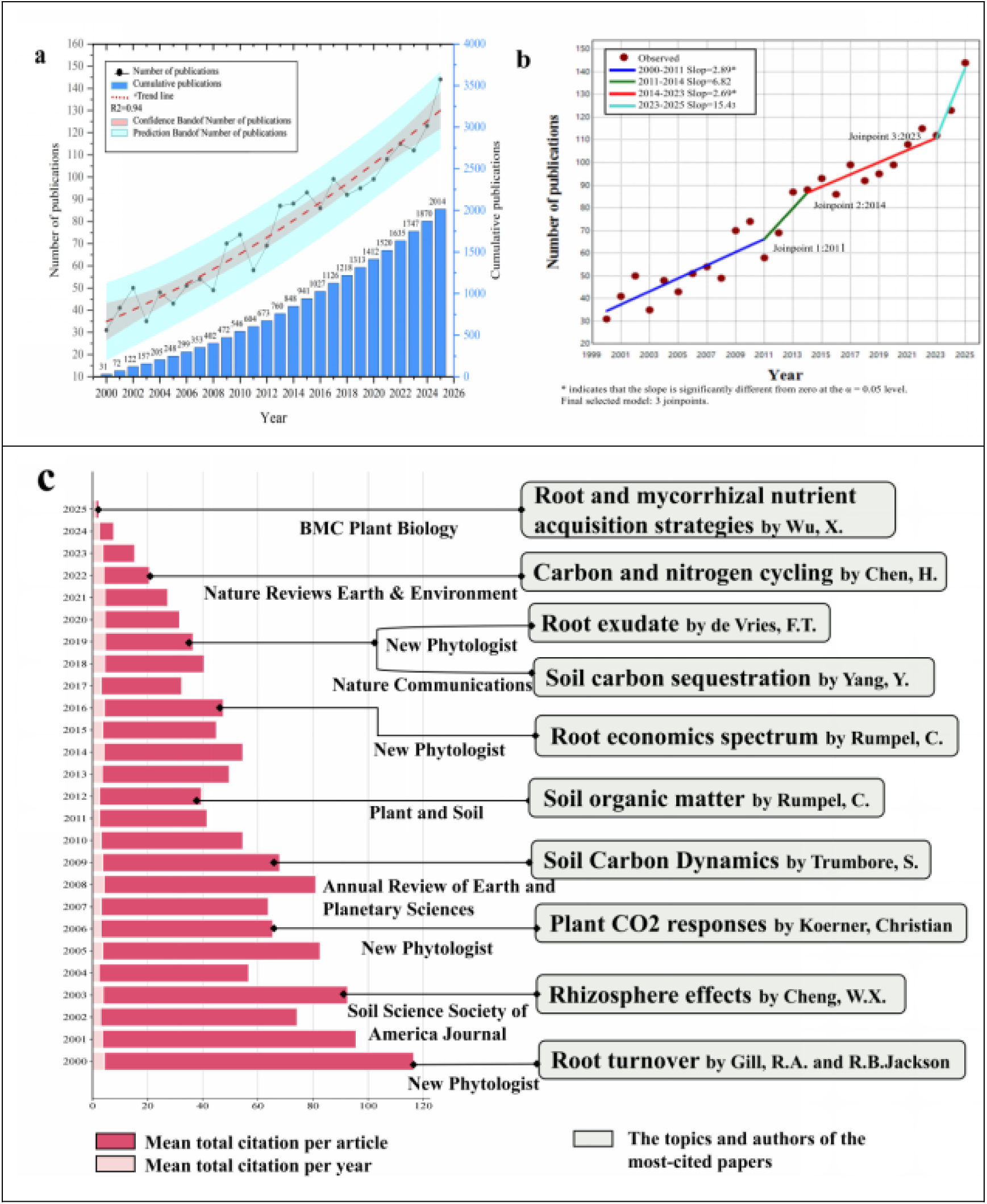
**The developmental trend of grassland root functional traits from 2000 to 2025 and the theme of the most highly cited article**. **a** Publication output distribution and trends over time. The red dashed line represents the trend line. **b** Phases of publication output in grassland root functional traits research. Asterisk indicates that the slope is significantly different from zero at the α = 0.05 level. Final selected model: 3 joinpoints.**c** The Evolution of themes of the most-cited articles.

### 2.2 Distribution of Publications by Country and Institution

An analysis of the contributing countries and institutions (based on author affiliations) reveals that the research output is dominated by a few key countries. In particular, China, the United States, and Germany have collectively propelled the development of this field, both in quantity of publications and in citations, reflecting their strong research programs in ecology and soil science. Figure 3a visualizes the global distribution of grassland root trait research output. Europe, North America, and Asia account for more than 85% of the global publication output. China has the highest number of publications, indicating intensive research efforts; the United States and Australia also make outstanding contributions, and together these three countries account for a large proportion of the total literature. International collaboration networks, illustrated in Figure 3b, show that while there are collaboration links (e.g., between China and the USA, and among European countries), the global cooperation network is not yet tightly integrated. Many research groups operate at the national or regional level, and cross-continental collaborations are relatively limited. Strengthening international cooperation could help form a more cohesive global research network. At the institutional level, As depicted in Figure 3c, Chinese research institutions such as the Chinese Academy of Sciences, the University of Chinese Academy of Sciences, and Northwest A&F University occupy central positions in the collaboration network of the top 100 institutions ranked by publication output, and are closely connected with other institutions.Chinese research institutions exhibit high output, with the Chinese Academy of Sciences being the single most prolific institution (Table 2). Other top institutions include the University of the Chinese Academy of Sciences, Northwest A&F University, Lanzhou University, and China Agricultural University in China; The University of Western Australia; and the University of Minnesota and the USDA in the United States. Notably, some institutions with moderate publication counts have very high citation averages (e.g., University of Minnesota, USDA), indicating that their contributions, while fewer in number, are particularly impactful. Overall, the data suggest that research activity is concentrated in a relatively small number of institutions, many of which are in China or have strong programs in plant and soil science.

**Figure 3.**
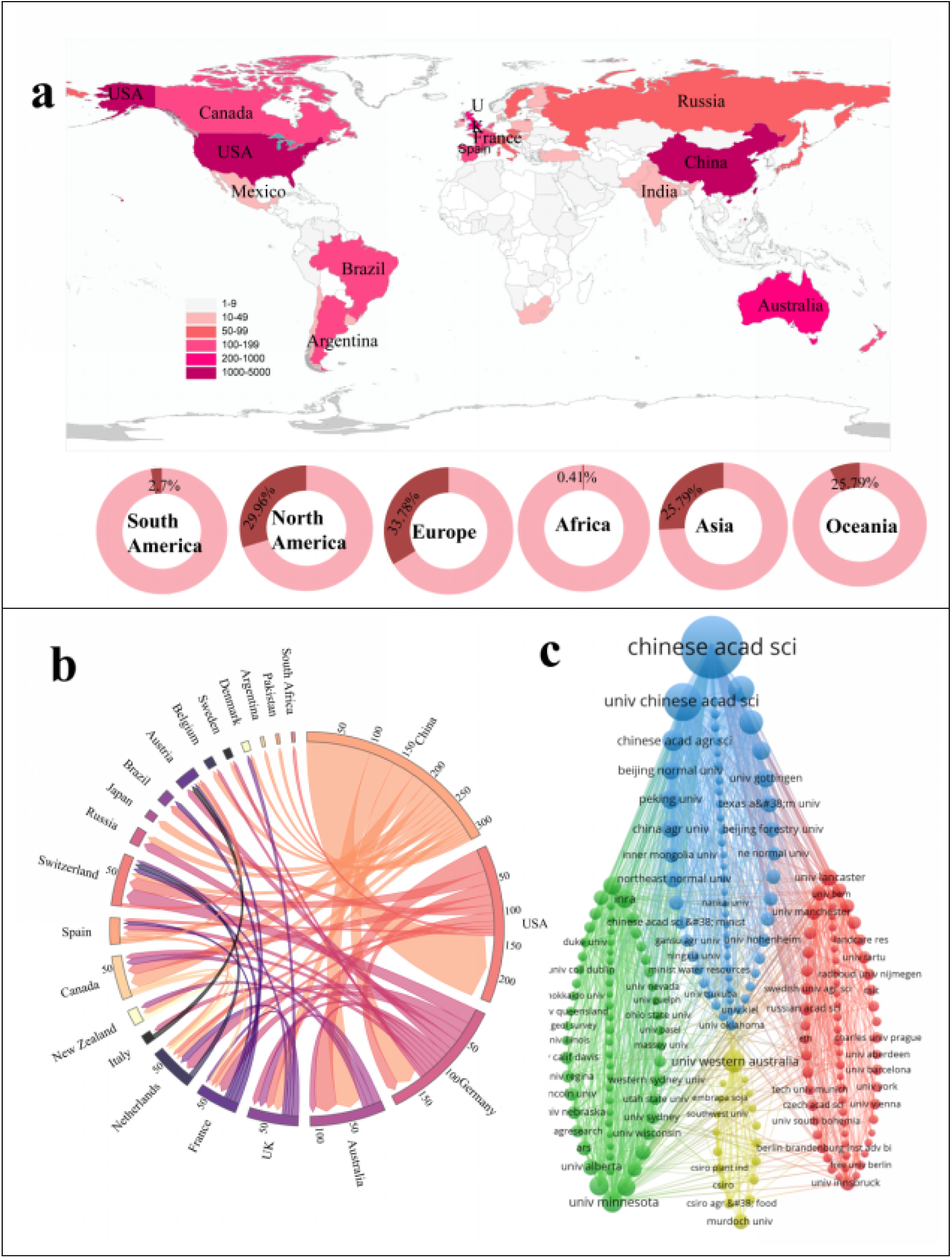
Country and Institution Analysis. a: World map showing the output of grassland root trait research by country (2000–2025). b: International collaboration network among major publishing countries (node size denotes publication count; line thickness indicates collaboration strength). c: Institutional collaboration network of major research institutions (with ≥100 publications), illustrating inter-institutional co-authorship ties.

**Table 2:** Citation counts and average citations per article for the top 10 research institutions (2000–2025).

| Rank | Organization | Articles | Citations | Mean TC per Art |
| --- | --- | --- | --- | --- |
| 1 | Chinese Academy of Sciences | 348 | 10435 | 30.0 |
| 2 | University of the Chinese Academy of Sciences | 128 | 3003 | 23.5 |
| 3 | Northwest A&F University | 60 | 1529 | 25.5 |
| 4 | Lanzhou University | 53 | 1359 | 25.6 |
| 5 | The University of Western Australia | 46 | 1427 | 31.0 |
| 6 | Chinese Academy of Agricultural Sciences | 41 | 533 | 13.0 |
| 7 | University of Minnesota | 41 | 4490 | 109.5 |
| 8 | United States Department of Agriculture (Usda) | 34 | 1976 | 58.1 |
| 9 | Peking University | 32 | 1430 | 44.7 |
| 10 | China Agricultural University | 31 | 1114 | 35.9 |

Note: “Articles” = number of publications; “Citations” = cumulative citations of those publications; “Mean TC per Art” = mean citations per paper, indicating average impact

### 2.3 Major Journals and Most Influential Papers

Analyzing the academic journals that publish research on grassland root functional traits, as well as identifying highly cited papers, helps in understanding the main channels of knowledge dissemination and the foundational literature of this field. The results show that the literature is spread across a range of journals in ecology, plant science, and soil science. Figure 4a demonstrates that Bradford’s Law can be applied, identifying a core set of journals contributing a large portion of the research output. These core journals (shaded region in the figure) include top-tier outlets such as Plant and Soil, Soil Biology & Biochemistry, Agriculture, Ecosystems & Environment, and Journal of Ecology, which have the highest number of publications on this topic and occupy central positions in the journal co-citation network. Figure 4b presents a journal co-citation network map (based on the top 100 journals by co-citation frequency). In this network, each node represents a journal; node size corresponds to how frequently that journal is cited in the grassland root trait literature, and links between nodes indicate how often two journals are cited together (reflecting related subject areas or communities). The clustering of journals in this co-citation map reveals disciplinary overlaps: for example, ecology and soil science journals cluster together, indicating an interdisciplinary knowledge base for this research field. Figure 4c also illustrates this point. In addition to journal analysis, identifying the most influential papers provides insight into key topics(Table 3). As noted, the most cited paper in the field is Rumpel et al. (2011) on soil carbon dynamics. Other highly cited works include studies by Freschet et al. on root trait effects on ecosystem functioning (2021), Bardgett et al. on linking above- and belowground ecology (2005), and Poirier et al. on root traits and soil organic matter stabilization (2018). These seminal papers have shaped current understanding and are frequently used as foundational references. The presence of such highly cited literature underlines certain enduring research themes—such as carbon cycling, nutrient cycling, and root morphology-function relationships—that form the knowledge base for grassland root functional trait research.

**Figure 4.**
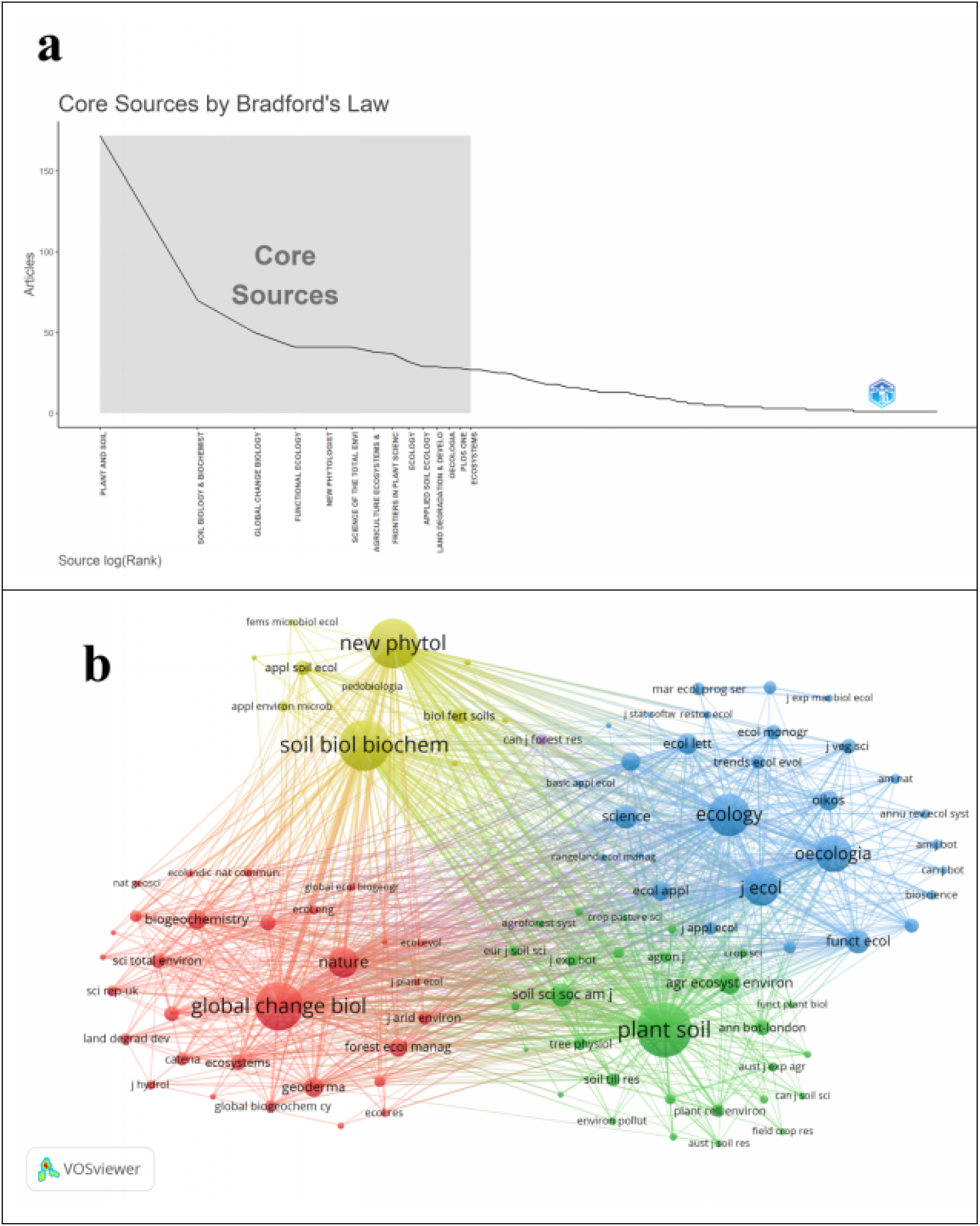
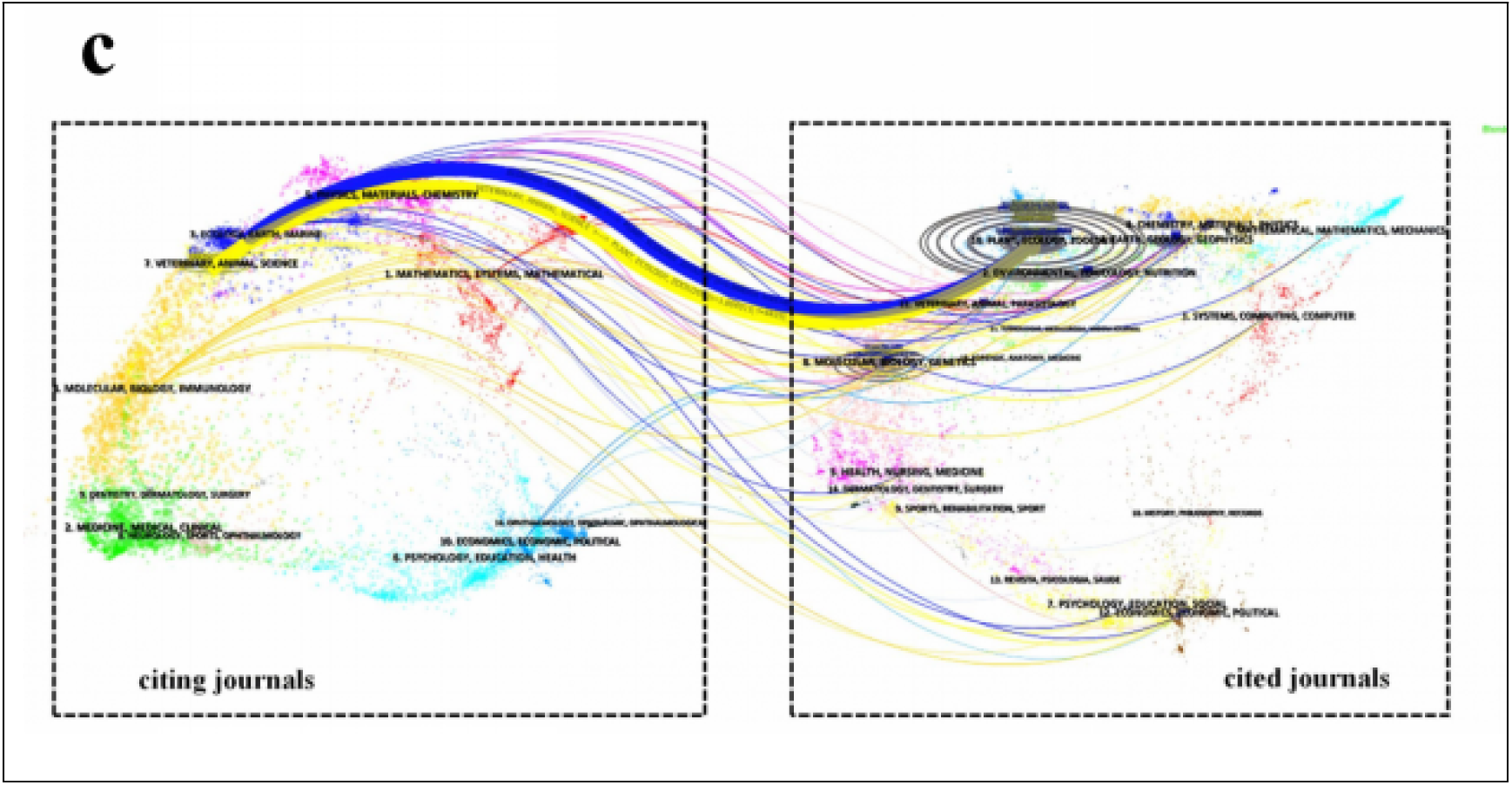
Journal Analysis. a: Bradford’s Law applied to journals in this field (core journal zone shaded). b: Co-citation network of the top 100 journals (node size represents citation frequency; line thickness represents co-citation strength). c: Dual-map overlay of journals publishing grassland root trait research, showing citation flows between fields

**Table 3:**
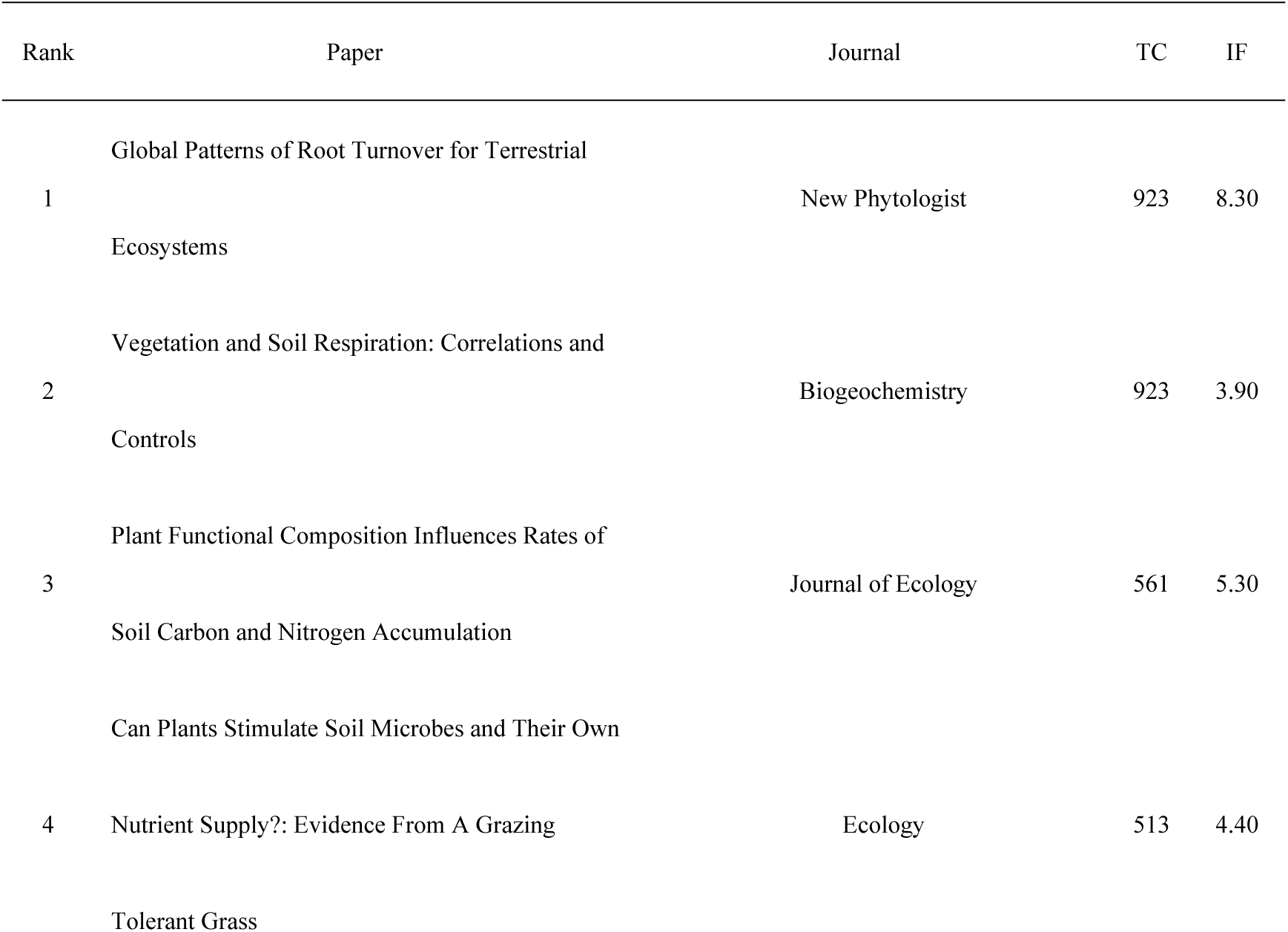
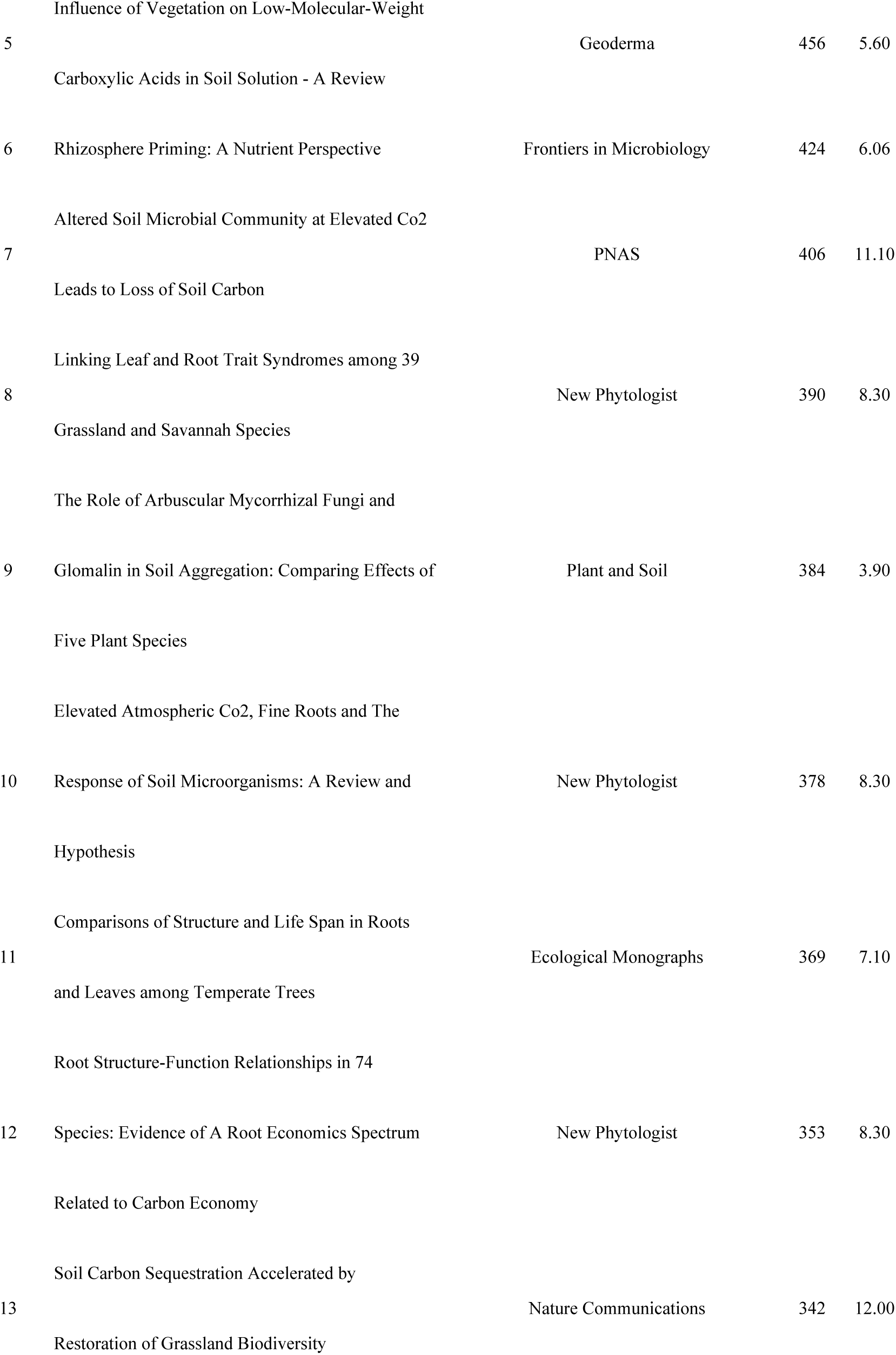
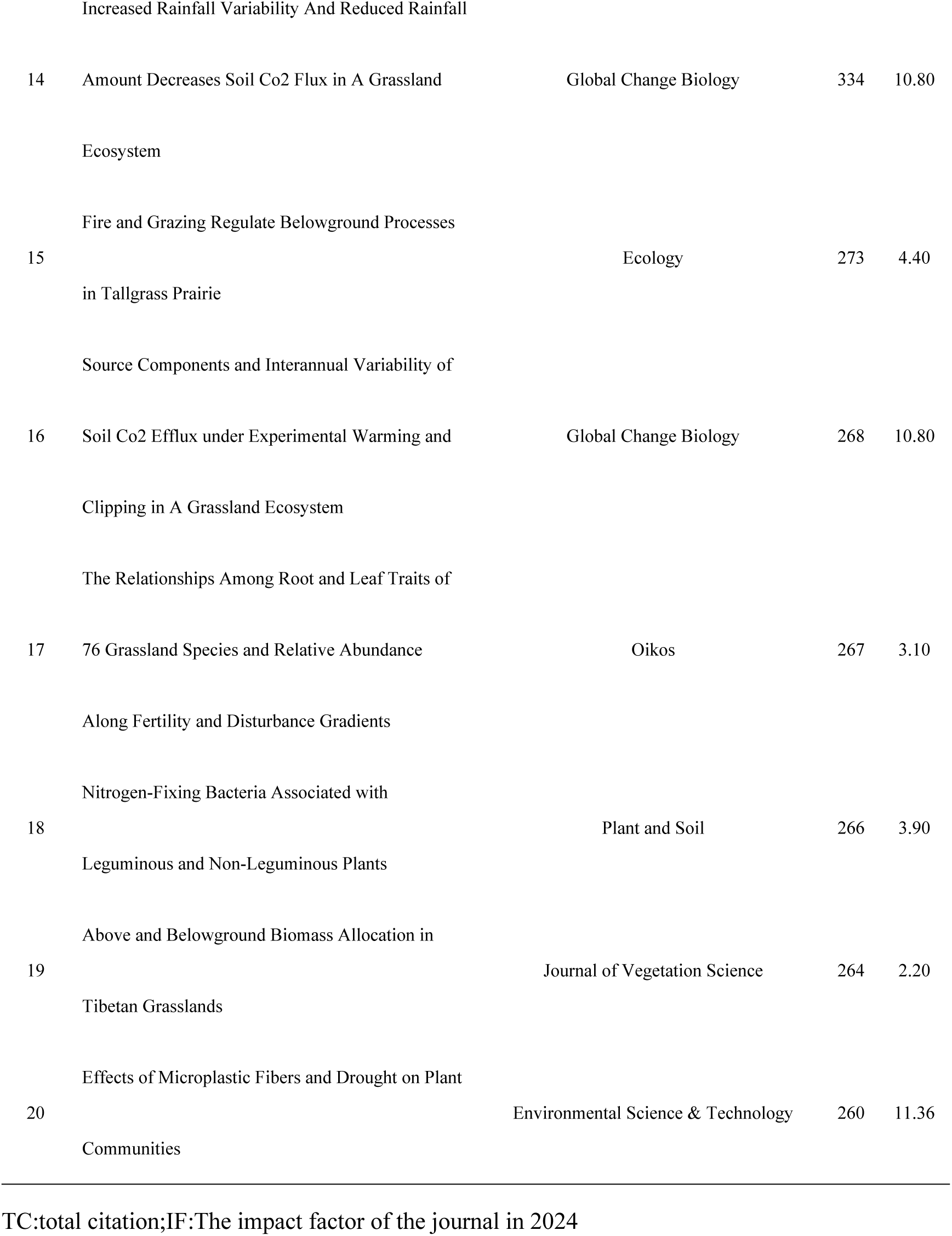
Top 20 Most Cited Articles on Grassland Root Functional Traits.

### 2.4 Author and Co-citation Analysis

Author-level analyses were conducted to identify the most prolific researchers and the intellectual structure of the field through co-citation patterns. The top 10 most productive authors and the top 10 most frequently co-cited authors are listed in Table 4. Notably, researchers such as Hans Lambers and Shuli Niu are among the most productive, each with 15–16 publications on grassland root traits, reflecting their active contributions to this area. In terms of influence, authors like Michael Bahn, Peter B. Reich, Gerd Gleixner, Liesje Mommer, David Tilman, and Yakov Kuzyakov feature prominently among the most co-cited, indicating that their work is frequently used as a knowledge base by other researchers. Figure 5a shows the co-authorship network of authors, which illustrates collaboration relationships among researchers who publish in this field. This network reveals several collaboration clusters, often corresponding to research groups or labs, sometimes centered around leading scientists. For instance, one cluster might consist of European researchers focusing on soil nutrient cycling and plant–soil interactions, while another cluster is centered on experts investigating root ecology under global change scenarios.

**Figure 5.**
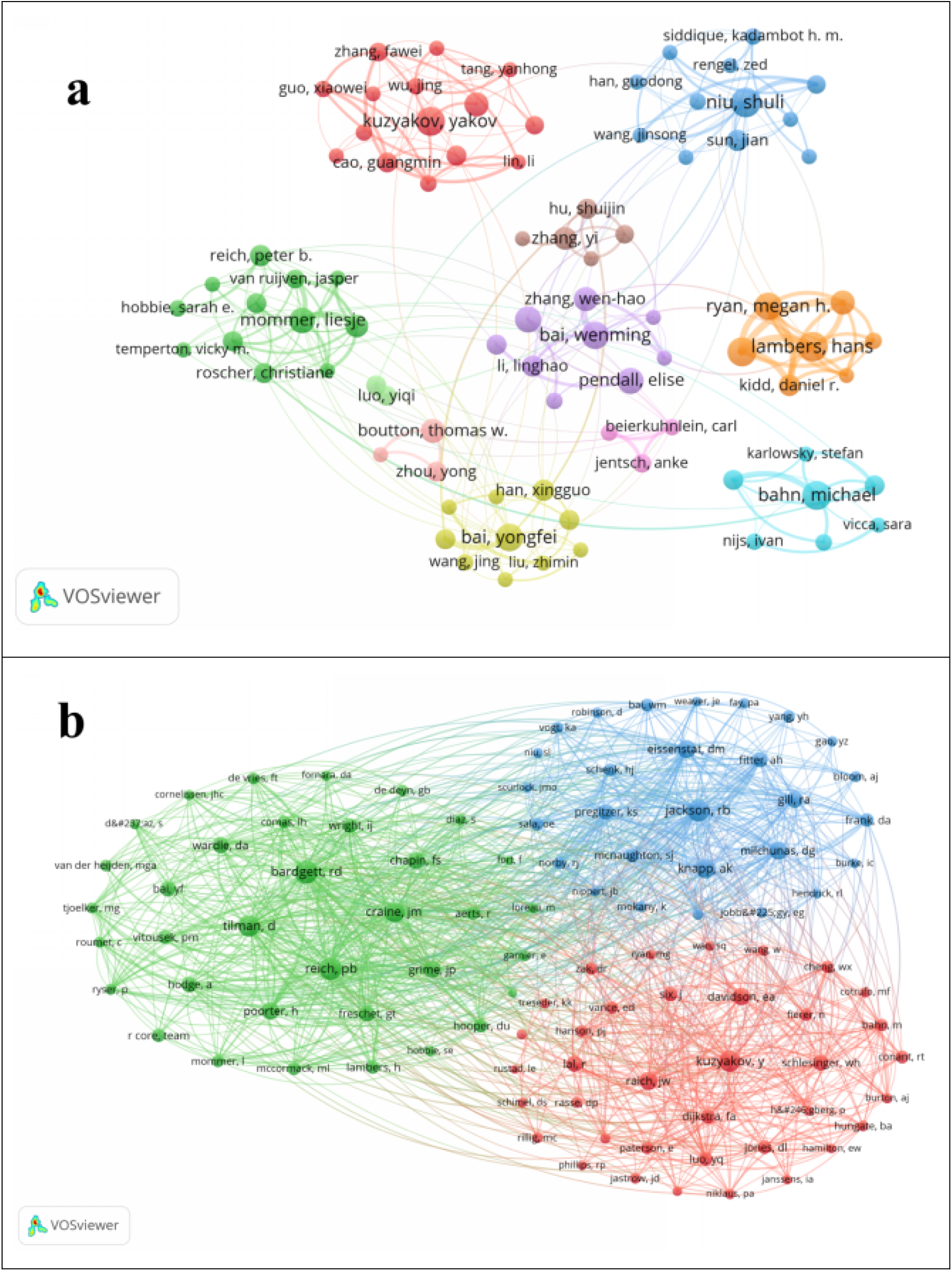
Author and Co-citation Analysis: a:The co-authors’ map of the research; b: In the co-cited authors’ map of the research

**Table 4:** The 10 Most Productive Authors and the Top 10 Most Co-cited Authors in the Study of Grassland Root Functional Traits.

| Rank | Author | Documents | Citations | TLS | Author | Documents | Citations | TLS |
| --- | --- | --- | --- | --- | --- | --- | --- | --- |
| 1 | lambers, hans | 16 | 503 | 54 | bahn, michael | 15 | 1270 | 25 |
| 2 | niu, shuli | 16 | 769 | 39 | reich, peter b. | 9 | 1065 | 17 |
| 3 | bahn, michael | 15 | 1270 | 25 | gleixner, gerd | 8 | 940 | 17 |
| 4 | kuzyakov, yakov | 15 | 647 | 11 | mommer, liesje | 12 | 912 | 40 |
| 5 | simpson, richard j. | 15 | 389 | 49 | reich, pb | 6 | 890 | 8 |
| 6 | bai, wenming | 14 | 607 | 29 | hobbie, sarah e. | 6 | 879 | 7 |
| 7 | bai, yongfei | 14 | 831 | 23 | tilman, david | 5 | 850 | 12 |
| 8 | ryan, megan h. | 14 | 323 | 43 | craine, jm | 5 | 838 | 10 |
| 9 | dijkstra, feike a. | 13 | 669 | 15 | bai, yongfei | 14 | 831 | 23 |
| 10 | pendall, elise | 13 | 682 | 10 | hasibeder, roland | 8 | 826 | 20 |

Figure 5b shows the co-citation network of authors (i.e., how often authors are cited together in the literature). This co-citation analysis provides a lens into the field’s intellectual structure. The co-citation network can be partitioned into a few major clusters of authors who tend to be cited together, suggesting they share related research foci or conceptual frameworks. For example, one prominent co-citation cluster (often highlighted as the red cluster in visualization) includes scholars like Yakov Kuzyakov and E.A. Davidson, who are well known for work on carbon and nitrogen dynamics in plant–soil systems. Another cluster (e.g., green cluster) is composed of authors such as R.D. Bardgett and D. Tilman, representing research on ecological effects of global changes (like climate change, biodiversity) on grassland ecosystems. A blue cluster might include experts such as R.B. Jackson and D.G. Milchunas, focusing on rhizosphere interactions and grassland management. These clusters reflect that the field encompasses multiple sub-disciplines, including soil biogeochemistry, ecosystem ecology, and global change biology, all converging on the theme of grassland roots.

Overall, the author collaboration and co-citation analyses indicate that while there are distinct research groups and thematic clusters, there is also considerable interconnection. This suggests a maturing research field with a growing community of scholars building upon each other’s work. Strengthening collaboration among these groups could further integrate the field and spur novel insights at the intersections of these clusters

### 2.5 Analysis of Research Hotspots and Evolutionary Trends

#### 2.5.1 Thematic Keyword Map and Evolutionary Trends of Research Hotspots

Keywords provide a concise summary of research content and are essential for identifying research directions and hotspots. An analysis of keyword frequencies (Table 5 and Figure 5a) shows the prominence of certain terms in the grassland root trait literature. High-frequency keywords correspond to topics that have received substantial attention and research investment, indicating mature or popular areas, whereas low-frequency keywords point to comparatively under-explored topics. For instance, keywords like “root biomass”, “root respiration”, “grassland”, “soil organic matter”, “climate change”, “grazing” and “root traits” appear with high frequency. This highlights that a considerable portion of research has focused on root biomass production, respiratory processes, and their relation to soil carbon dynamics and climate or land-use factors. The presence of “nitrogen” and “drought” as notable keywords underscores the interest in nutrient cycling (particularly nitrogen addition/deposition effects) and plant responses to water limitation in grasslands. Meanwhile, terms like “arbuscular mycorrhizal fungi (AMF)” and “rhizosphere” also feature prominently, reflecting a significant research focus on root–microbe interactions and belowground ecological processes. A keyword co-occurrence network was constructed to visualize how topics group into clusters of related research (Figure 5b). In this network, each node is a keyword (with node size proportional to its frequency), and links between nodes represent co-occurrence in the same publications. The network reveals six major clusters (hotspot themes) of co-occurring keywords, effectively partitioning the field into thematic areas: Cluster 1 (Red) – represented by keywords such as “root biomass”, “climate change”,“grazing”, “root functional trait” and “pasture”. This cluster primarily focuses on how environmental factors like climate warming and grazing disturbance affect plant root biomass and functional traits in grassland ecosystems. It also explores the bidirectional interactions between plant roots and grassland ecosystem processes under these stresses. The prominence of both climate-related terms and grazing suggests an emphasis on global change factors and land-use in grassland root research (e.g., how drought or warming plus grazing pressure influence root growth and grassland functioning)[54–58]. Cluster 2 (Green) – includes keywords such as “nitrogen,” “nitrogen addition,” “restoration”, “grass”, “Leymus chinensis” (a common grass species of the Eurasian steppe), “Tibetan Plateau” and “root/shoot ratio”. This cluster is centered on nutrient enrichment (especially nitrogen deposition/addition experiments) and grassland restoration studies, often in alpine or temperate grasslands (e.g. on the Qinghai-Tibetan Plateau). Research in this cluster examines how nitrogen inputs affect ecosystem structure and function, plant traits (including root:shoot allocation), and strategies for degraded grassland restoration[59, 60]. Cluster 3 (Blue) – dominated by terms like “arbuscular mycorrhizal fungi (AMF)”, “CO₂”, “root exudation”,“rhizosphere” and “carbon sequestration”. This cluster focuses on soil microbial interactions, particularly the role of AMF and root exudates in carbon cycling. It emphasizes mechanisms by which AMF symbiosis and rhizosphere processes contribute to carbon sequestration in grassland ecosystems. For example, studies in this cluster investigate how AMF facilitate the transfer and storage of carbon in soil and how root exudates influence soil carbon decomposition or stabilization[61, 62]. Cluster 4 (Yellow) – characterized by keywords like “grassland”, “soil organic matter”, “root turnover” and “carbon cycling”. This cluster focuses on plant diversity and ecosystem function, examining how plant functional diversity (especially belowground) impacts grassland productivity and soil organic matter dynamics. A key theme is fine root turnover as a driver of soil carbon and nutrient cycling. Studies often explore how root litter inputs and turnover rates under different plant diversity scenarios contribute to soil organic carbon pools and nutrient availability[63]. Cluster 5 (Purple) – includes terms such as “root,” “carbon,” “soil,” “land use change,” “soil properties,” and “carbon allocation”. This cluster is mainly about how land-use changes (e.g., conversion between grassland, cropland, pasture) and management practices affect soil properties and soil organic matter through changes in root traits and carbon allocation. It addresses issues like organic carbon and nitrogen turnover in soils when grasslands undergo land-use transitions, highlighting the role of root systems in mediating soil fertility and structure under land-use change[64–66]. Cluster 6 (Cyan) – features keywords related to soil factors and root physiology, such as “root respiration,” “soil microbial biomass,” “soil properties,” and “fine root” (often appearing in context with nutrient cycling and stress responses). This cluster underlines the integration of soil microbial processes with root functional studies. For example, it includes topics like how soil microbial biomass influences fine root branching intensity, or how root physiological processes (respiration, nutrient uptake) interact with soil conditions. The clustering of these terms indicates a holistic approach to studying root traits in the context of soil ecosystems—bridging root physiology and soil microbiology[67, 68].In terms of thematic maturity(figure 6c), the landscape of research keywords remained relatively sparse and fragmented during the early period (2000–2014), with most terms appearing in cooler hues (blue to purple), indicating low or sporadic usage. A clear inflection point emerged around 2015, when the number of active keywords began to gradually increase, signaling the consolidation and expansion of the field.

**Figure 6.**
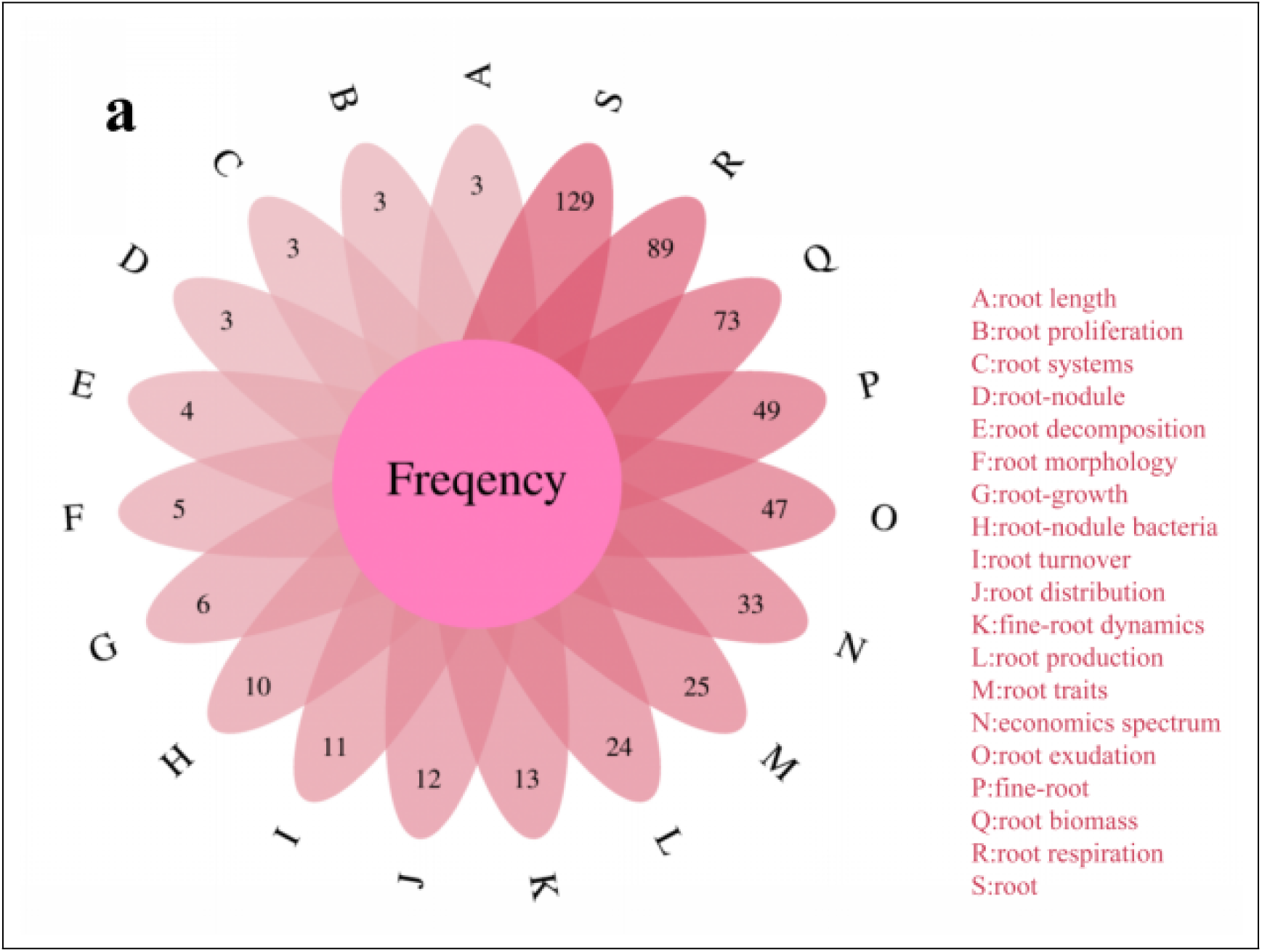
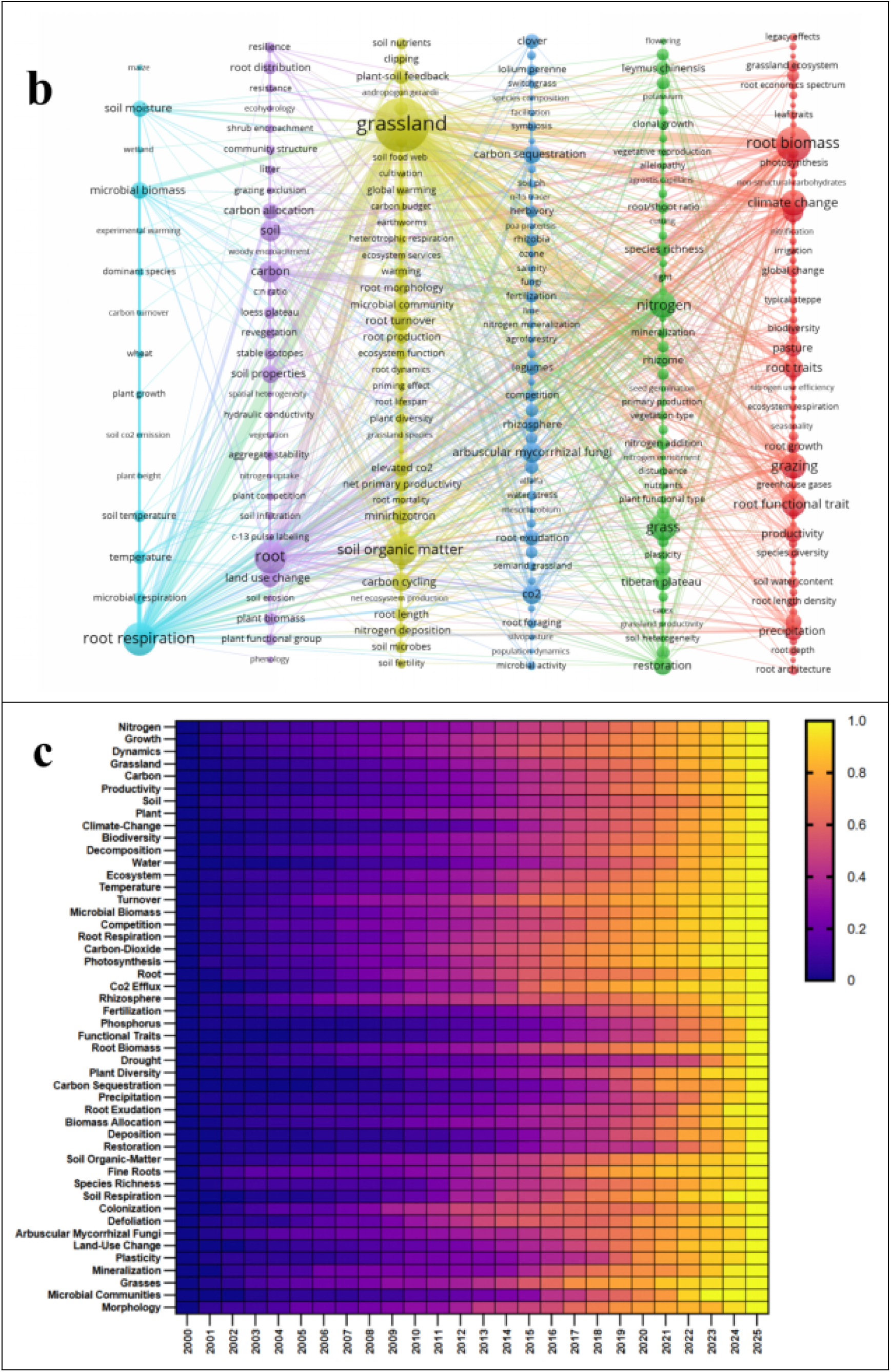
a:Keyword Analysis:a Frequency of keywords related to root functional traits;b :Cooccurrence network knowledge domain map of keywords (frequency ≥ 5) ; c :Temporal heatmap of high-frequency keywords.

**Table 5:** Frequency of Appearance and Association Strength of the Top 25 Keywords.

| Rank | Keyword | Cluster | Occurrences | Total Link Strength |
| --- | --- | --- | --- | --- |
| 1 | grassland | 4 | 372 | 900 |
| 2 | root biomass | 1 | 156 | 358 |
| 3 | root respiration | 6 | 140 | 378 |
| 4 | root | 5 | 133 | 334 |
| 5 | soil organic matter | 4 | 131 | 317 |
| 6 | grass | 2 | 95 | 171 |
| 7 | nitrogen | 2 | 91 | 301 |
| 8 | grazing | 1 | 87 | 202 |
| 9 | climate change | 1 | 85 | 249 |
| 10 | root functional trait | 1 | 76 | 204 |
| 11 | carbon | 5 | 59 | 177 |
| 12 | soil | 5 | 58 | 129 |
| 13 | root traits | 1 | 54 | 157 |
| 14 | arbuscular mycorrhizal fungi | 3 | 50 | 114 |
| 15 | drought | 1 | 50 | 150 |
| 16 | CO<sub>2</sub> | 3 | 48 | 150 |
| 17 | root turnover | 4 | 48 | 144 |
| 18 | restoration | 2 | 46 | 112 |
| 19 | carbon cycling | 4 | 44 | 129 |
| 20 | land use change | 5 | 43 | 110 |

Overall, the keyword co-occurrence analysis paints a picture of a field that is divided into these thematic areas yet interconnected. The identified clusters correspond closely to the six research directions highlighted earlier. Each cluster represents an active hotspot of research, ranging from climate change impacts to nutrient cycling and restoration practices. Importantly, the distribution of keywords across clusters suggests that grassland root trait research is inherently interdisciplinary, encompassing climate science, soil science, plant physiology, and ecology.

#### 2.5.2 Identifying Research Frontiers

To identify emerging research frontiers, we analyzed temporal keyword trends and “burst” keywords (keywords with a sudden increase in attention). The temporal analysis of keyword usage provides another perspective on how research emphases have shifted over time (Figure 7a). In the early 2000s, keywords related to basic ecological processes (e.g., “soil organic carbon,” “nutrient cycling,” “root respiration”) and fundamental grassland terms (“grass,” “roots”) were dominant, reflecting foundational studies establishing baseline understanding of grassland root ecology. Starting around 2010–2012, terms such as “grassland,” “climate change,” “grazing,” and “root biomass” became more prominent, indicating an expansion of scope to include ecosystem responses and management-related factors. In the past five years (approximately 2018–2024), there has been a notable rise in terms like “root traits,” “root functional trait,” “fine root,” and topics like “nitrogen addition/deposition,” signifying a shift toward more refined questions about specific root trait functions and anthropogenic factors. This trend reflects a growing emphasis on the functional role of roots (rather than just root quantity) and on experimentally manipulating factors like nutrient deposition to probe ecosystem responses. The evolution of research themes can be summarized in three stages: (1) a “Basic Ecological Understanding” stage, focusing on characterizing fundamental root and soil processes in grasslands (e.g., root contributions to soil carbon); (2) a “Physiological Mechanism Exploration” stage, delving into physiological and micro-scale processes (such as root respiration, exudation, and rhizosphere interactions), often with advanced techniques to investigate root traits at finer scales; (3) an “ Application-Oriented Research”stage, concentrating on environmental changes (climate, nitrogen deposition, etc.) and stress responses, including ecological restoration strategies.The keyword burst analysis (Figure 7b) highlights terms that have experienced sharp, short-term surges in usage, pointing to emerging or rapidly expanding research frontiers. The top five keywords with the highest burst strength in recent years are “loess plateau,” “root traits,” “tallgrass prairie,” “biomass allocation,” and “nitrogen addition”. Notably, “tallgrass prairie,” “lolium perenne,” and “plants” exhibit the most sustained burst durations, signifying a long-standing research focus on specific grassland ecosystems (tallgrass prairie), dominant pasture species (Lolium perenne), and foundational plant ecological processes.Several keywords that first appeared earlier in the timeline—such as “carbon dioxide,” “root respiration,” and “soil organic carbon”—have witnessed renewed or substantially heightened attention in recent years. Over the last five years, entirely new research focal points have also emerged, including “plant functional traits” (expanding beyond root-centric traits to integrate aboveground and whole-plant functional strategies), “drought,” and “strategy” research. The rise of “drought” reflects growing concern over climate change-driven water stress and its ecological consequences, while the increased focus on “strategy” indicates a shift toward investigating plant adaptive mechanisms in dynamic environmental conditions. Combining the co-occurrence clusters and the temporal analyses, we see a clear trend: The field has broadened and deepened, with early work establishing fundamental linkages between roots and ecosystem processes, mid-phase research integrating climate and land-use factors, and recent work honing in on functional trait mechanisms and applications for ecosystem management. In summary, grassland root trait research is transitioning into a more integrative stage, addressing complex, multi-factor scenarios and bridging fundamental science with practical management solutions. This reflects a shift from purely academic exploration toward research that can inform adaptive management of grassland ecosystems under global change.

**Figure 7.**
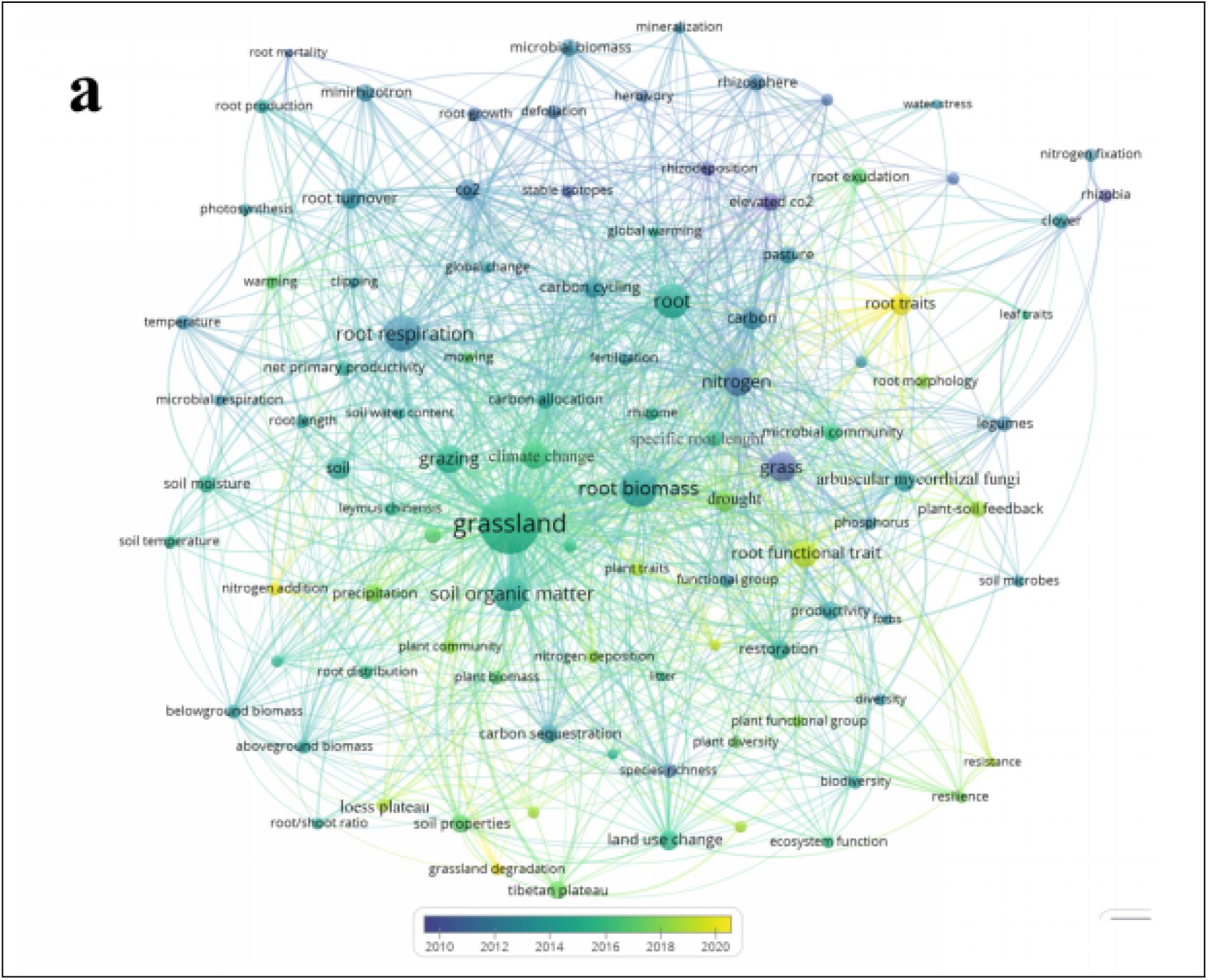
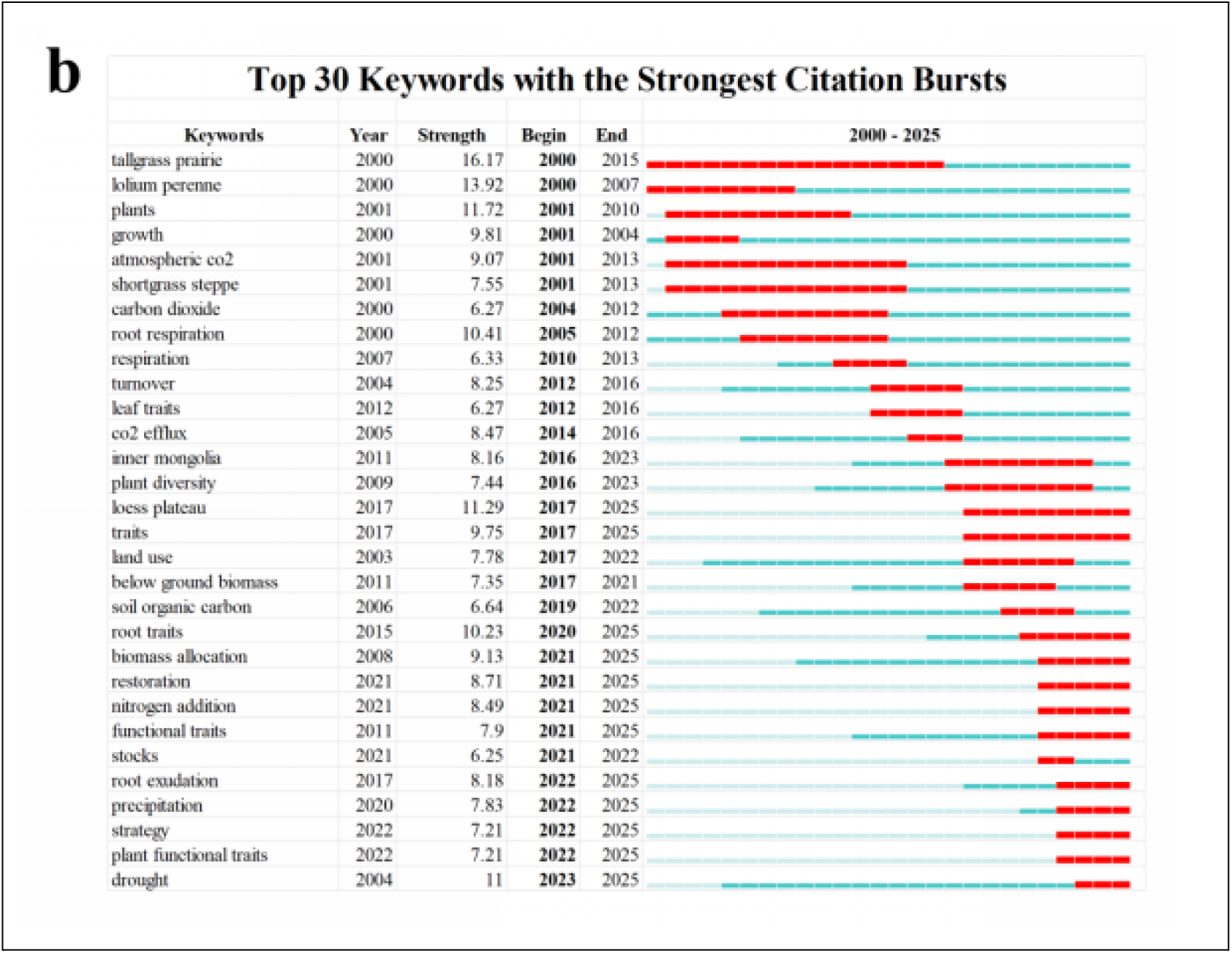
Identifying Research Frontiers:a Time Trend Chart of Keywords.b Visualization of the Top 30 Keywords with the Strongest Citation Bursts

## 3. Discussion

### 3.1 Multidimensional Characteristic Studies of Grassland Root Functional Traits

One prominent frontier in this field is the study of the multidimensional nature of root functional traits[69, 70]. Early in trait ecology, the concept of a single-axis “Root Economic Spectrum” (RES) was proposed to summarize trade-offs in root traits[71]. However, accumulating evidence indicates that grassland plant root traits possess multiple dimensions of variation rather than fitting neatly into one fast–slow continuum. This multidimensionality spans various aspects of root form and function, including morphology, physiology, and ecological roles of roots. The absorptive function of roots is often determined by a suite of interrelated traits (such as total root biomass, root diameter, SRL, etc.), and the interplay among these traits affects resource acquisition efficiency[9, 72]. For instance, a plant’s ability to acquire soil resources depends not on a single trait but on a combination—e.g., a fine-rooted grass with high SRL may explore soil volume efficiently, but if those roots are too thin they may have shorter lifespan or lower mycorrhizal colonization, introducing trade-offs[73]. Additionally, root trait multidimensionality is evident in anatomical structures and geometry. Allometric relationships between root cortical and stele diameters, and independent variation in cell wall thickness and tissue density, contribute extra complexity to root trait space. These multiple axes of variation mean that a single index or spectrum is insufficient to capture plant strategies; rather, roots simultaneously vary along several functional axes (e.g., an axis of resource acquisition vs. conservation, an axis of root tissue construction costs, and an axis of root symbiotic dependency, etc.). Importantly, the multidimensionality of root traits influences not only how plants acquire resources but also their stress tolerance and contributions to ecosystem functions. For example, a grassland species might adapt to nutrient-poor conditions by developing finer roots with high SRL (maximizing exploration) but also increasing root tissue density (to enhance longevity under stress)—combining traits that span different dimensions. Different environmental drivers can act on different trait dimensions. Climate factors (like temperature and precipitation) and soil factors (like nutrient availability, texture) each play distinct roles in shaping fine-root trait variation, often leading to complex, multidimensional gradients of root trait responses across environments. Recent research confirms that no single environmental gradient explains all root trait variation; instead, multiple factors each pull root traits in different directions, reinforcing the idea of multiple trait axes. Recognizing this, some scholars have proposed conceptual frameworks and models for multidimensional root trait space. For instance, Zhao et al. (2020) constructed a model of multidimensional fine-root trait space, providing new insights into belowground trait variation by integrating morphological, chemical, and architectural traits[72]. Experimental work also supports multidimensional trait thinking: Ding et al. examined how climatic factors (especially precipitation) and soil nutrients shape root system attributes, finding that changes in precipitation regime can simultaneously affect multiple root trait dimensions (morphology, chemistry, etc.) by altering resource heterogeneity[74]. Similarly, Kong et al.’s study of tropical tree fine roots revealed significant intra-root variation in carbon composition at the molecular level, which in turn affects where those roots fall in an economic spectrum, adding a molecular-scale dimension to root trait variability[75]. Such findings introduce new perspectives on root diversity, showing that even within “fine roots” there is chemical and functional differentiation that influences their role in ecosystems. Despite progress, methods and tools to quantify root trait multidimensionality remain underdeveloped, and a unified standard for measuring and comparing multidimensional root traits is lacking. This has become a bottleneck for advancing this research direction. There is a clear need for standardized protocols and integrative measurement approaches (e.g., combining root imaging, chemical analysis, and functional assays) to accurately capture the multiple dimensions of root traits across species and systems. We expect that research on root trait multidimensionality will continue to grow, with efforts directed at developing these methods and analytical tools. By doing so, researchers can more comprehensively reveal the multidimensional functional characteristics of grassland root systems and better understand their ecological significance. Our bibliometric analysis supports the rising importance of this topic: keywords and clusters related to multidimensional trait perspectives are emerging as hotspots. Concepts like “root trait variation,” “trait spectrum,” and multi-trait analyses are increasingly common in recent literature. In summary, exploring the multidimensional characteristics of grassland plant root functional traits is crucial for deepening our understanding of how belowground parts contribute to plant adaptation and ecosystem function. This line of inquiry will provide a more nuanced framework that goes beyond simple spectra, ultimately improving predictions of plant behavior and ecosystem processes under varying environmental conditions.

### 3.2 Interactions between Grassland Root Functional Traits and Climate Change

Climate change—encompassing increased atmospheric CO₂, rising temperatures, altered precipitation regimes, and more frequent extreme weather events—has profound impacts on natural ecosystems and poses urgent challenges for sustainable management. In grassland ecosystems, multiple elements of climate change can directly or indirectly influence plant carbon allocation strategies and absorptive root traits, thereby regulating key ecosystem functions[76]. Understanding how grassland root functional traits respond to climate change factors is critical for predicting ecosystem feedbacks and resilience[77]. Elevated CO₂ (eCO₂) and increased temperatures (warming) are two of the most studied climate drivers in this context. These factors can significantly alter root biomass allocation and root function. Under eCO₂, for example, plants often increase their fine root biomass, extend root length, and grow roots to greater depth, effectively “foraging” deeper for water resources[78]. Such responses enhance water acquisition under CO₂-driven increased growth, but they also change the vertical distribution of carbon inputs into the soil. Warming similarly tends to prompt plants to allocate more biomass belowground. Experiments have shown that in warmer conditions, grasses and other plants will invest in deeper root systems—figuratively “rooting deeper”—to access moisture from deeper soil layers as surface soils dry out[76]. This strategy of deeper rooting under warming helps plants cope with surface drying, but it comes with a potential “double-edged sword” effect: on one hand, more carbon is transported to and stored in deeper soil layers (which could enhance soil carbon sequestration); on the other hand, stimulating decomposition in deep soil (the so-called “carbon bomb” in permafrost or long-stored carbon) could be activated, leading to release of CO₂ from soil that had long-term storage. Different plant functional groups in grasslands (e.g., grasses vs. forbs vs. legumes) exhibit varying root trait responses to climate stressors[79]. Under drought stress—a component of climate change that many grasslands are experiencing with greater frequency—grasses (family Poaceae) commonly adapt by increasing root diameter and root tissue density. Thicker, denser roots can be more resistant to drought-induced cavitation and can explore soil more thoroughly for scarce water, albeit often at the cost of reduced specific root length. Other functional groups, such as forbs, may adopt different drought strategies (e.g., enhancing taproot depth or association with mycorrhizae)[80].These differences underscore that root trait plasticity under climate stress is highly context- and species-dependent, and they emphasize the key role of root traits in plant adaptation to environmental changes. Changes in root traits under climate stressors have cascading effects on soil microbial processes and biogeochemical cycling. For instance, a shift to larger diameter, lower SRL roots under drought can alter the quality and decomposability of root litter inputs to soil[81]. Research has found that drought-induced changes in plant traits (including roots) can modify litter decomposition rates, either strengthening or weakening the feedback of vegetation to climate change[82]. One study noted that while drought might lead to more recalcitrant litter (slowing decomposition), the plasticity of root and shoot traits does not always translate into proportional changes in decomposition, indicating a complex interplay and highlighting that root trait changes can have non-linear effects on ecosystem carbon feedbacks[83]. Overall, climate change influences on root traits—by altering carbon inputs and nutrient cycling rates—have significant indirect effects on grassland carbon balances and nutrient dynamics. Our keyword co-occurrence analysis confirms that “climate change” forms a distinct hotspot cluster of research in this field. Terms related to warming, CO₂, drought, and associated disturbances (like “grazing” which often interacts with climate effects) are clustered, indicating intensive study of these factors. The red cluster identified earlier aligns with this, highlighting that the community is keenly interested in how environmental changes affect root traits and, reciprocally, how those trait changes feedback to ecosystem processes. This body of work collectively explores the bidirectional feedbacks between plant roots and ecosystems under climate stress: for example, warming and CO₂ affecting root growth and turnover, which in turn affect soil carbon release or storage.

Despite progress, a notable bottleneck remains a lack of comprehensive understanding of root trait response mechanisms under combined climate factors. Most experiments to date have studied single-factor effects (e.g., warming alone, or drought alone). However, real-world climate change involves simultaneous changes (e.g., warmer and drier conditions, or increased CO₂ with altered precipitation). There is a relative scarcity of multifactor climate manipulation experiments that consider, for instance, warming + altered rainfall + elevated CO₂ all together. Such experiments are complex but necessary to reveal potential interactions (synergistic or antagonistic) in root trait responses. For example, increased CO₂ might mitigate some negative effects of drought on root growth (through enhanced carbohydrate supply), or warming might exacerbate drought impacts—these scenarios need testing. Future research trends in this area will likely focus on more scenario-based, long-term experiments that simulate complex climate futures. Long-term field trials combining factors (e.g., open-top chambers or Free-Air CO₂ Enrichment plus rainfall shelters for drought) can elucidate how adaptive strategies of root traits manifest under realistic, multifactor global change conditions. Concurrently, there is growing emphasis on incorporating root trait parameters into ecosystem and Earth system models. Traditionally, many ecosystem models simplified or ignored root trait variation, but as data accumulate, models are beginning to include traits like SRL, root depth distribution, and root turnover rates to improve predictions of carbon and water fluxes under climate change. By integrating functional root traits into models, we can achieve better forecasts of grassland responses to climate change and more robust assessments of carbon sequestration potential and ecosystem service maintenance in grassland regions. In summary, climate change significantly influences grassland root functional traits, and these trait responses have important implications for ecosystem dynamics. Ongoing and future research—through multi-factor experiments and trait-informed modeling—is crucial to unravel the complex response mechanisms and feedback loops, thereby improving our ability to predict and manage grassland ecosystems in a changing climate.

### 3.3 Synergistic Interactions between Grassland Root Functional Traits and Soil Microorganisms

The interactions between plant roots and soil microorganisms (especially mutualistic fungi and rhizosphere bacteria) represent a key dimension of grassland ecology and are an important frontier in root trait research. Grassland plant roots do not function in isolation; they form symbiotic and associative relationships with myriad soil microbes that can significantly influence nutrient uptake, plant health, and soil processes[84]. Among these, the symbiosis with arbuscular mycorrhizal fungi (AMF) is particularly prominent, as most grassland plant species (notably grasses and legumes) form AMF associations. Research into root–microbe interactions has thus become a hot topic within the study of root functional traits[85].

One fundamental finding is that plant root traits can drive the abundance and community structure of associated soil microorganisms. For example, there is a strong relationship between root morphological traits and AMF colonization: plant species with larger root diameters tend to support higher AMF colonization rates and greater fungal biomass in roots, whereas species with very high SRL (thin roots) often show lower AMF colonization[86]. This suggests an inverse relationship where extremely fine, high-SRL roots might rely less on mycorrhiza (perhaps because they explore soil so extensively themselves), whereas thicker roots benefit from fungal partners to extend their foraging range. In essence, root morphology can determine, to some extent, the degree of symbiosis: roots are not just passive recipients of fungi; their traits actively influence which and how many fungi colonize them.

The ecological functions conferred by AMF are considerable. AMF are well known to enhance host plant nutrient acquisition, particularly for relatively immobile nutrients like phosphorus (P). By extending the effective reach of the root system through their hyphal networks, AMF allow plants to tap phosphorus beyond the immediate root zone. In doing so, they expand plant nutrient acquisition pathways and can improve plant nutrition and growth[87]. Moreover, AMF and other root-associated microbes play crucial roles in soil carbon dynamics. Through root exudates and the growth of fungal hyphae, they influence soil microbial community structure and the balance of soil organic matter decomposition vs. stabilization. For instance, AMF presence can alter the composition of rhizosphere microbial communities, sometimes stimulating decomposition of organic matter (a process known as the “priming effect”) or aiding in soil carbon stabilization by contributing to aggregate formation and physical protection of carbon in soil[88]. AMF hyphae also transport plant-derived carbon into the soil matrix over a broader volume, effectively distributing carbon inputs and potentially enhancing soil carbon sequestration in grasslands.

Studies under stress conditions further highlight the value of root–microbe synergy. Under adverse conditions such as salinity or nutrient deficiency, AMF can improve host plant performance by various means. For example, under salt stress, AMF colonization has been shown to boost the host plant’s photosynthetic efficiency and even leaf thickness, thereby improving plant stress resistance[89]. By enhancing plant water and nutrient status and modulating hormonal signals, mycorrhizae help mitigate the negative impacts of abiotic stress on plants[90, 91]. This demonstrates that mycorrhizal symbiosis significantly contributes to plant tolerance of harsh environments—a mechanism by which grassland ecosystems maintain functional diversity and stability in the face of stress. Despite these advances, many of the mechanistic underpinnings of root–microbe interactions remain only partly understood. The plant–AMF symbiosis involves complex signal exchanges and resource trading: plants supply carbohydrates to fungi, and fungi supply nutrients (and sometimes water) to plants[92]. The molecular dialogues (signaling molecules for colonization, coordination of nutrient transfer) are subjects of active research, and many details are still unclear. For instance, how plants regulate carbon allocation to roots vs. to mycorrhizae, how they “decide” when the symbiosis is beneficial vs. not, and how different microbial partners might be favored under different conditions—these are questions requiring further study. Additionally, beyond AMF, grassland roots interact with a myriad of other microbes: nitrogen-fixing bacteria, phosphorus-solubilizing bacteria, pathogens, etc. The networks of interaction (the root–microbiome system) are complex, and current research is only beginning to unravel how diverse microbial communities influence root trait expression and vice versa[93].

On a community and ecosystem level, different plant species exhibit distinct strategies in their root–microbe interactions, which in turn can influence community composition. Comparative studies have shown, that fine-rooted grass species often have a different “mycorrhizal strategy” than coarse-rooted species. Fine-rooted species (with high SRL and extensive branching) often rely more on direct uptake (with less mycorrhizal dependency), typically showing lower AMF colonization, exuding fewer organic acids, and exhibiting lower rhizosphere phosphatase activity. Under low soil P, these fine-root strategists strongly increase their own root branching and length to acquire nutrients[94]. In contrast, coarse-rooted species invest in the symbiotic route: they tend to have higher AMF colonization, and their roots secrete more compounds that can mobilize soil P (perhaps because thicker roots can’t physically explore soil as extensively, so they rely on fungi and chemistry)[95, 96]. These contrasting strategies illustrate a sort of complementarity or trade-off between “root-alone” vs. “root-plus-microbe” nutrient acquisition strategies. Interestingly, soil microbial biomass itself can feedback to influence root architecture[97, 98]. Some studies indicate that the abundance of soil microbes can affect fine root branching intensity, suggesting that even subtle changes in microbial community size or composition can modify root developmental patterns[99]. Minor alterations in root structure (like changes in branching) can, in turn, directly impact soil microbial habitats and communities, potentially creating feedback loops.

Different types of mycorrhizal associations (e.g., arbuscular vs. ectomycorrhizal) also lead to different plant responses to environmental change. Grasslands are predominantly AMF-associated, but understanding from other systems (forests with ectomycorrhizae) suggests that mycorrhizal type can influence plant strategies under warming or nutrient enrichment[100]. For instance, ectomycorrhizal plants may respond differently to warming than AMF plants in terms of biomass allocation. While grasslands mostly involve AMF, this raises interesting questions about whether different AMF strains or other endophytes might analogously lead to varied outcomes.

In sum, research on the synergistic interactions between grassland root functional traits and soil microorganisms has made significant strides. We now appreciate that root–microbe interactions operate on multiple levels: macro-level community patterns (which species associate with which microbes under what conditions) and micro-level physiological and biochemical mechanisms (how nutrients and signals are exchanged) have both been explored in depth[101]. Bibliometric analysis shows that work centered on arbuscular mycorrhizae forms an independent cluster of hotspots (the blue cluster), underscoring the pivotal role of AMF in grassland root ecology. Another identified cluster (cyan) focuses on soil microbial factors like microbial biomass in relation to root respiration and soil properties, indicating an inseparable link between soil microbial processes and root trait studies. However, a current challenge is the lack of a comprehensive understanding of the complex interactions between diverse soil microbial communities and the equally diverse plant root traits. For example, how do multiple microbial groups (bacteria, fungi, others) interact with a suite of root traits simultaneously? What are the overall effects on ecosystem function when considering entire microbiomes rather than single symbionts? These questions remain open. There is also a need to integrate root–microbe interactions into ecosystem models and management practices — for instance, using microbial inoculants to alter root trait expression (as discussed in applications below) or predicting how shifts in soil microbiomes (due to climate or land use) will affect plant trait-mediated processes. Addressing these will require interdisciplinary approaches, combining microbiology, plant physiology, and ecology. As techniques like high-throughput sequencing (for microbiomes) and in situ root phenotyping improve, we can expect rapid progress. The synergistic interplay between roots and soil microbes is fundamental to grassland resilience and productivity; thus, unraveling these connections will not only advance theory but also guide practices for grassland restoration and sustainable management (e.g., through microbial amendments or breeding for root traits that foster beneficial microbes).

### 3.4 Response of Grassland Root Functional Traits to Land-Use Change

Land-use change—such as conversion between native grassland, cropland, pasture, and the intensification or abandonment of grazing—has a profound impact on grassland ecosystems, often comparable in magnitude to climate change effects[102, 103]. Grassland root functional traits respond sensitively to land-use and management changes, and these responses can alter ecosystem processes like productivity and soil carbon storage[104, 105]. Understanding these responses is crucial for managing grasslands sustainably amidst agricultural expansion and shifting land-use practices.

One major effect of land-use change in grasslands is on rooting depth and distribution[106]. For example, when natural grasslands are converted to croplands or heavily grazed pastures, there is often a shift in plant community composition towards species with shallower or thinner roots (e.g., annuals or grazing-tolerant grasses) at the expense of deep-rooted perennials. This can lead to reduced overall root depth and biomass in deeper soil layers, affecting the soil’s ability to sequester carbon at depth and access deep water or nutrients[107]. Conversely, restoration of degraded grasslands or reduction of grazing pressure can allow re-establishment of deep-rooted perennial grasses and forbs, thereby increasing root inputs to deeper soil and improving soil structure[108].

Land-use change also typically impacts root trait syndromes related to the plant economic spectrum. Intensive agriculture or overgrazing tends to favor fast-growing species with “acquisitive” trait profiles (high SRL, low RTD, short-lived roots) which rapidly capture nutrients when available, whereas undisturbed or lightly managed grasslands often maintain more “conservative” root traits (thicker, denser roots with longer lifespan)[109, 110]. These shifts have implications: acquisitive roots might promote quick nutrient cycling (faster turnover), while conservative roots contribute to long-term carbon storage in soil[111]. Indeed, land-use driven changes in root turnover rates can be a major driver of differences in soil organic carbon between managed and natural systems[112]. Grazing is a particularly important land use in grasslands. Moderate grazing can sometimes stimulate root growth (through compensatory growth and enhanced nutrient cycling from manure), but heavy grazing generally reduces root biomass, especially at depth, as plants allocate more to re-growing shoots and less to maintaining extensive root systems[113]. Grazing can also change species composition (favoring grazing-resistant or unpalatable species), which often means a shift in dominant root traits of the community. For instance, grazing-resistant species might have higher root reserves (carbohydrates in roots) to rapidly regrow after defoliation, which is a functional trait shift relevant to ecosystem recovery[114].

Our analysis of keyword clusters indicates that land-use change appears as a key term (notably in the purple cluster identified in the co-occurrence analysis, which includes “land use change,” “soil properties,” and “carbon allocation”)[115]. This shows that many studies link root traits with changes in land management and observe effects on soil properties and carbon cycling[116]. Indeed, research has demonstrated that changes in land use can alter root-driven processes such as soil aggregation (through changes in root exudates and fungal associations) and soil carbon input/output balance[117]. For example, conversion of grassland to cropland often results in a reduction of root-derived carbon inputs and a disturbance of soil structure, leading to declines in soil carbon stocks[118]. On the other hand, practices like grassland restoration, reforestation with silvopasture, or introduction of deep-rooted species aim to reverse these effects by enhancing root inputs and improving belowground traits conducive to carbon sequestration[119].

A current bottleneck in this research direction is the lack of long-term, systematic monitoring of how different land-use practices affect belowground processes, particularly in deeper soil layers and over decadal timescales. Most studies are short-term or focus on surface soil and immediate root responses. We have an incomplete understanding of how chronic land-use pressures (like continuous overgrazing or repetitive tillage) gradually change root trait distributions and soil function. Also, cross-site comparisons are limited: grasslands in different climatic zones (savannas vs. steppes vs. prairies) may respond differently to the same land-use due to baseline differences in dominant species and trait syndromes, which introduces uncertainty in generalizing model parameters for carbon cycling[120].

Future research needs to address these gaps by conducting more comparative studies across regions and ecosystem types, combined with long-term experiments (e.g., long-term pasture exclosure vs. grazed plots, or decades-long cropping vs. restored grassland trials). Such studies would reveal general patterns as well as region-specific responses of grassland root traits to land-use change. Additionally, integrating ecology with agronomy and soil science can help apply root trait knowledge to land management. For example, understanding root trait responses could guide rational grazing management (optimal stocking rates and rotation to maintain deep-rooted species), or fire management, as well as inform species selection for restoration (choosing plant species with root traits that improve soil structure and carbon storage). These insights aim to achieve win-win outcomes for production and ecological health, such as maintaining forage yield while also sequestering carbon and preventing soil degradation[121].

In summary, strengthening research on the linkages between grassland root functional traits and land-use changes is of great theoretical and practical significance. Theoretical, because it deepens our understanding of ecosystem stability and service provisioning under anthropogenic influence; practical, because it informs how we manage grasslands for sustainable use. Grassland management strategies that account for and leverage root trait dynamics—such as fostering deeper rooting through moderate grazing or mixed-species swards—are likely to enhance both productivity and soil conservation. Given the global extent of grasslands and the pressures they face, this knowledge is vital for achieving long-term ecological sustainability and climate mitigation goals in these ecosystems.

### 3.5 Application of Grassland Root Functional Traits in Agriculture and Ecological Restoration

Applying the insights from grassland root functional trait research to agricultural production and ecological restoration is a promising direction that has gained momentum in recent years[122]. Bridging the gap between fundamental research and practical management, this area emphasizes using knowledge of root traits to improve grassland sustainability, enhance crop and forage productivity, and restore degraded lands. Our analysis also indicates that research related to “application practice” is heating up, with an increasing number of studies focusing on how manipulating or selecting for certain root traits can yield practical benefits. For example, in the co-occurrence analysis, the green cluster (involving keywords like “nitrogen addition” and “restoration”) reflects efforts to use root trait knowledge to guide management experiments in alpine grasslands and restoration strategies.

Current specific research directions in this applied arena include the following aspects: Firstly, optimizing root structure and function of crops and forage species to enhance tolerance to abiotic stresses (e.g., drought, salinity)[123]. This involves breeding or selecting varieties with desirable root traits or adopting cultivation practices that encourage deeper and more extensive root systems. For instance, breeding programs are increasingly looking at root traits (like deeper roots or higher root length density) as targets for improving drought resistance in forage grasses and cereals. A deeper or more robust root system can improve water uptake under drought and access nutrients from a larger soil volume, thereby stabilizing yields in stress-prone environments[124]. Similarly, in saline soils, selecting plants with roots that exclude or compartmentalize salt effectively can improve survival. By manipulating planting density or soil conditions (like subsoiling or bio-drilling with cover crops), farmers can also encourage crops to develop more extensive root networks.

Secondly, adjusting grassland management practices (e.g., fertilization, pesticide use, mowing frequency) to balance belowground and aboveground biomass allocation[125, 126].The goal is to enhance carbon sequestration and productivity simultaneously. For instance, rational fertilizer use can promote root growth rather than excessive shoot growth, leading to more carbon input belowground. Similarly, altering mowing or grazing frequency can allow plants enough recovery time to rebuild root biomass, improving soil organic matter input and long-term productivity[127]. Studies have shown that moderate fertilization and controlled grazing, as opposed to intensive management, can lead to better root growth and soil carbon accumulation. A well-managed grassland will allocate biomass in a way that supports both yield (forage for livestock or hay) and soil health (root-derived carbon)[128]. Thus, developing guidelines for fertilization or mowing that explicitly consider root responses (not just immediate aboveground yield) is a key application of root trait knowledge.

Thirdly, using plant growth-promoting rhizobacteria (PGPR) or other beneficial microbiota to alter plant root traits for improved crop performance. Soil inoculants containing beneficial bacteria or fungi can stimulate root development and modify root architecture[129]. For example, certain rhizobacteria are known to increase root branching or root hair length, thereby enhancing a plant’s nutrient and water uptake capacity[130].They can also improve soil structure by secreting polysaccharides that bind soil particles (enhancing aeration and water retention). Practice has shown that such biological interventions can reduce reliance on chemical inputs and lead to more resilient agro-ecosystems. Incorporating PGPR inoculants into grassland management or crop cultivation (especially in organic or sustainable systems) leverages natural processes to achieve healthier root systems and, consequently, better plant growth and soil quality.

Fourthly, directly targeting soil carbon sequestration through management that encourages beneficial root traits[131]. For instance, increasing the proportion of deep-rooted perennial species in a pasture mix can drive more carbon into deeper soil horizons, where it is more likely to be stored long-term. Practices such as silvopasture (integrating deep-rooted trees with pasture) or periodic sowing of cover crops with extensive root systems are being explored as ways to improve root-mediated carbon inputs. Additionally, reducing disturbance (like no-till or reduced tillage in grassland–crop rotations) helps maintain root networks and soil structure, thereby enhancing carbon sequestration. By managing for root traits—like promoting root systems that turn over slowly or produce more root litter—land managers can influence the soil carbon sink capacity of grasslands.

These applied research directions demonstrate how a trait-based understanding can translate into concrete techniques for improving grassland and agricultural outcomes. In agricultural grasslands, the aim is often a win-win: achieving high productivity (for livestock or biomass) while also maintaining or improving soil health and carbon stocks[132]. For ecological restoration, the emphasis might be on selecting species or management regimes that rebuild soil structure, prevent erosion, and restore nutrient cycling—roles largely mediated by root systems[133]. Our literature review indicates that these topics—drought-resistant roots, balanced fertilization for roots, PGPR effects, etc.—are gaining traction. For example, studies have documented that introducing legumes with deep roots into a degraded grassland can accelerate restoration of soil fertility, or that inoculating seedlings with mycorrhizal fungi speeds up establishment in restoration projects. In practice, integrating these insights requires interdisciplinary collaboration among plant breeders, agronomists, soil scientists, and ecologists. It also requires field experimentation to tailor general principles to local conditions. However, the trend is clear: grassland management and restoration strategies are increasingly root-centric, recognizing that sustainable solutions lie as much belowground as above. By focusing on root functional traits, practitioners can improve outcomes such as drought resilience, reduced fertilizer needs, enhanced soil carbon sequestration, and long-term productivity.

### 3.6 Coupling of Root Functional Traits with Grassland Ecosystem Functions

Building on the above applications, an emerging research frontier is the development of technologies and methodologies specifically aimed at regulating or enhancing root functions in agricultural ecosystems. This area takes a more technological and innovation-driven approach to root trait management, seeking to optimize root system performance to improve crop yield and sustainability. It is considered one of the potential future hotspots for grassland and agroecosystem research, as indicated by increasing references to “root phenomics” and “root engineering” in recent literature. To address current limitations in root research and application, several breakthrough directions have been proposed:

a. Developing Multimodal Root Phenomics Technologies and Standardized Observation Systems: This involves creating advanced techniques (imaging, sensors, tomography) to measure root traits non-destructively and in real time, and establishing standardized protocols for data collection and analysis. For instance, improved root imaging systems (e.g., X-ray CT scanning, ground-penetrating radar, rhizotrons with image analysis) can capture root architecture and growth dynamics with high resolution[134]. Standardizing these measurements across studies will enable better comparison of root traits and more rapid breeding for root characteristics. By innovating in root phenotyping (analogous to aboveground high-throughput phenotyping), researchers and breeders can more efficiently select for desirable root traits like deeper rooting, higher root length density, or more root hairs[135, 136].
b. Designing Long-Term Experiments with Multiple Environmental Factor Interactions: This direction calls for experimental setups that simultaneously consider various environmental factors (drought, nutrient availability, elevated CO₂, warming, etc.) to better mimic complex real-world conditions[137, 138]. The goal is to understand root trait responses under multifactor scenarios and identify traits or trait combinations that confer resilience in such contexts. For example, a long-term field experiment might impose both drought and nutrient stress to see which root traits help plants cope with the combined stress—data that can inform breeding and management. By integrating multiple factors, these experiments will improve prediction models and help disentangle trait responses in highly variable environments[139, 140].
c. Elucidating Metabolic Interaction Networks and Functional Regulation Pathways between Roots and the Microbiome: This direction focuses on understanding the intricate metabolic exchanges and signaling pathways between plant roots and soil microorganisms[141]. It aims to uncover how roots and microbes co-regulate processes such as nutrient uptake, growth, and stress responses. Technologies like metagenomics, metabolomics, and stable isotope tracing are employed to map nutrient flows and signal transduction in root–microbe networks[142]. A better grasp of these interaction networks could lead to novel ways of managing or engineering microbiomes to benefit root function (for example, encouraging microbes that induce roots to grow deeper or branch more under certain conditions). This fundamental understanding is crucial for any bioinoculant or microbial management strategy (as discussed in 3.4) to be effective and reliable[143, 144].
d. Constructing Integrated Multiscale Models for Cross-Level Prediction of Root Trait Dynamics: This direction aims to develop models that can predict root trait dynamics across different scales—from individual root segments to whole plants, and from plot-level to ecosystem-scale. These models would integrate data from molecular, physiological, and ecological levels to forecast how root systems will develop under various scenarios (climate change, management regimes, etc.) and how those root dynamics will translate to ecosystem outcomes (yield, carbon sequestration, etc.)[145]. Achieving this requires merging fine-scale process knowledge (like root respiration in response to nitrogen, hormone-driven root growth responses, etc.) with larger-scale factors (soil heterogeneity, competition in communities). If successful, such models could become powerful decision-support tools for agriculture and conservation, allowing stakeholders to simulate how altering a root trait (through breeding or management) might impact long-term soil health and productivity.

By systematically advancing these research and innovation directions, it is hoped that current bottlenecks—such as the difficulty of observing roots in situ over time, the separation of temporal and spatial scales in experiments, and the fragmented understanding of root mechanisms—can be overcome. Collectively, these efforts promise both theoretical innovations (deeper insight into plant–soil interactions) and practical solutions (better tools and strategies for grassland and crop management). For example, improved root phenotyping can directly aid breeding programs to produce the next generation of stress-resilient crops with root systems tailored to future climate conditions (deep roots for drought, prolific roots for nutrient uptake, etc.)[146]. Long-term multifactor experiments will ensure that those traits are effective under realistic conditions, rather than only in simplified scenarios. Understanding root–microbe metabolic networks might lead to custom “soil probiotics” that farmers can apply to fields to elicit beneficial root trait changes, reducing the need for fertilizers or irrigation. And integrated modeling can tie everything together, forecasting outcomes and guiding policy (e.g., selecting crop varieties for carbon farming initiatives based on modeled root trait contributions to soil carbon sequestration).

In conclusion, applying root functional trait knowledge is transitioning from observational science to a new phase of innovation and technology deployment in agroecosystems. By embracing these interdisciplinary and multiscale approaches, scientists and practitioners aim to harness root traits as levers for improving productivity, sustainability, and climate resilience in grassland and agricultural landscapes. Advancing these frontier areas will be key to overcoming current challenges and unlocking the full potential of root traits for adaptive management of grassland ecosystems under global change.

## 4. Conclusion

This study employed bibliometric approaches and knowledge mapping tools (CiteSpace, VOSviewer) to systematically assess global research developments in grassland root functional traits from 2000 to 2025, drawing on data from the Web of Science Core Collection. Our analysis demonstrates a marked and sustained growth in international research output, with China, the United States, and Germany emerging as prominent contributors. Notably, the Chinese Academy of Sciences has established itself as a leader in this field. However, despite these advances, transnational research collaboration remains limited, as indicated by relatively weak inter-institutional connectivity and restricted knowledge exchange across borders. Through multidimensional analysis, we identified six major research frontiers. First, multiscale root trait analysis has advanced our understanding of root morphological, physiological, and chemical diversity across organizational and spatial gradients. Second, research on climate–root trait interactions is elucidating how climatic drivers and root functional traits interact to influence ecosystem resilience to environmental change. Third, the study of root–microbe ecological networks is uncovering the vital roles that plant–soil microbial interactions play in soil health and nutrient cycling. Fourth, investigations into root plasticity under land-use change highlight the adaptive capacity of root traits to varying intensities of anthropogenic disturbance, with implications for ecosystem stability and restoration. Fifth, the links between root traits and ecosystem services—such as carbon sequestration, productivity, and nutrient cycling—are increasingly recognized as central to grassland function. Finally, the development of root optimization technologies in agriculture points to promising innovations for sustainable production systems. To address current research gaps, we propose four key directions: (1) advancing multimodal root phenomics and standardizing trait measurement protocols; (2) designing long-term, multifactorial experiments to capture complex environmental interactions; (3) elucidating metabolic pathways and regulatory networks between roots and the microbiome; and (4) constructing integrated, multiscale predictive models of root trait dynamics. Pursuing these directions will help overcome existing bottlenecks—such as limited integration across spatial and temporal scales and incomplete mechanistic understanding—and will provide the theoretical and practical foundation for adaptive grassland ecosystem management under global change.

## CRediT authorship contribution statement

Yongxue Feng: Conceptualization, methodology, formal analysis, writing - original draft preparation. Yang Gao: Editing. Junqin Li: Editing.Yuting Yang: Editing.Jing Yin: Editing. Shu Yang: Editing. Zhihua Jiang:Editing. Lili Zhao: Editing. Puchang Wang: Conceptualization, writing - review and editing, supervision, funding acquisition. Xiangtao Wang:Conceptualization, writing - review and editing, supervision, funding acquisition.

